# Coordinated pre- and postsynaptic protein dynamics underlie rapid Sema4D-mediated inhibitory synapse assembly

**DOI:** 10.64898/2026.01.21.700908

**Authors:** Zachary Pranske, Suzanne Paradis

## Abstract

In the mammalian hippocampus, synapses are either excitatory or inhibitory as defined by the presynaptic neurotransmitter (glutamate or GABA, respectively) and the specific ligand-gated ion channel receptors localized to the postsynaptic specialization. While numerous studies explore the formation of excitatory synapses, the process of inhibitory synapse formation is less understood. Using both loss- and gain-of-function approaches, our lab previously identified the class 4 Semaphorin Sema4D as a key regulator of inhibitory synaptogenesis. Here, using recombinant Sema4D protein as a tool to rapidly induce GABAergic synapse formation in cultured hippocampal neurons, we employ two-channel live imaging to identify changes to pre- and postsynaptic protein dynamics during inhibitory synapse formation. We find that Sema4D treatment promotes the mobility of presynaptic GAD65 protein assemblies while having a negligible effect on the behavior of the postsynaptic gephyrin scaffold, leading to increased colocalization of these proteins. In addition, Sema4D treatment promotes the recruitment of GABA_A_Rγ2 subunits to immature gephyrin scaffolds, suggesting that Sema4D primes these scaffolds for receptor recruitment. Surprisingly, we observe new colocalization events between existing gephyrin and GABA_A_R puncta, suggesting that clustering of either the gephyrin scaffold or the GABA_A_R is sufficient to nucleate assembly of the postsynaptic specialization. Overall our results support a model in which Sema4D signaling coordinates dynamic changes in both pre- and postsynaptic compartments to assemble inhibitory synapses on rapid timescales.

**Significance Statement:** The assembly of new synaptic contacts requires precise coordination of specialized proteins in pre- and postsynaptic neurons. Inhibitory synapses, which suppress neuronal activity and are essential for circuit stability, contain distinct molecular components, yet the mechanisms governing their assembly remain poorly understood. We used Sema4D, a protein that rapidly induces inhibitory synapse formation, as a molecular tool to dissect how synaptic proteins on either side of the synaptic cleft are coordinated in space and time. Using live imaging we show that Sema4D acts on both pre- and postsynaptic compartments to recruit synaptic proteins with spatiotemporal precision. Together, these findings define the sequence of molecular events underlying inhibitory synapse assembly and have implications for neurodevelopmental disorders in which inhibition is disrupted.

## Introduction

Synapses are the core unit of cell–cell communication in the nervous system. Early studies of excitatory, glutamatergic synapse assembly in hippocampus revealed that synapse formation begins with the establishment of a transient contact between an axon and a dendrite (Scheiffele, 2003; Ziv and Garner, 2004). Next, mobile, pre-clustered protein assemblies are localized to the presynaptic compartment (Ziv and Garner, 2004; McAllister, 2007), while postsynaptic maturation is marked by accumulation of neurotransmitter receptors and scaffolding proteins (Prange and Murphy, 2001; Bresler et al., 2004; Ziv and Garner, 2004). The action of signaling molecules such as LRRTMs, Teneurins, Neuroligin/Neurexins, and Semaphorin/Plexins is thought to be essential for both initial contact formation and subsequent recruitment of synaptic proteins (Kuzirian et al., 2013; McDermott et al., 2018; Südhof, 2018, 2021), although it remains broadly unclear at which stage(s) of synapse development these molecules act.

Proper regulation of inhibitory, GABAergic synapse assembly is essential for circuit function, but compared to glutamatergic synapse formation, only a handful of studies have directly addressed the steps involved using live imaging approaches (Wierenga et al., 2008; Dobie and Craig, 2011; Kuriu et al., 2012; Villa et al., 2016; Frias et al., 2019). Collectively these studies revealed coordinated mobility and gradual accumulation of proteins associated with GABAergic synapses at axon–dendrite crossings over several hours, but did not resolve molecular recruitment events on the order of seconds to minutes. This limitation is significant, as the earliest stages of synapse assembly are likely governed by rapid, transient molecular events. Thus, there remains a substantial gap in understanding how molecular signals acutely regulate cellular processes that transform nascent contacts into mature GABAergic synapses.

A major barrier to addressing this problem has been the lack of tools to manipulate inhibitory synapse formation with temporal precision. We discovered a rapid molecular trigger for inhibitory synapse formation: the extracellular domain of the transsynaptic signaling protein Semaphorin 4D (Sema4D) is sufficient to induce the formation of new inhibitory synapses between hippocampal neurons within 30 minutes (Kuzirian et al., 2013; McDermott et al., 2018; Adel et al., 2023). These synapses become functional within 2 hours as revealed by in vitro and ex vivo physiology and in vivo mouse models (Kuzirian et al., 2013; Acker et al., 2018; Adel et al., 2023). Both loss-of-function (Paradis et al., 2007; Raissi et al., 2013) and gain-of-function studies (Kuzirian et al., 2013; McDermott et al., 2018; Adel et al., 2023) demonstrate that Sema4D specifically regulates GABAergic synapse formation without affecting glutamatergic synapses, making it a uniquely selective and temporally precise tool for studying inhibitory synaptogenesis.

Beyond their developmental roles in axon guidance, neuronal migration, and tissue morphogenesis, Semaphorins and their receptors have emerged as critical mediators of synaptogenesis (Paradis et al., 2007; Joo et al., 2013; Duan et al., 2014; Koropouli and Kolodkin, 2014; Uesaka et al., 2014). Sema4D is a transmembrane protein that can be cleaved from the pre-or postsynaptic membrane, allowing its extracellular domain to signal either in cis or in trans to Plexin-B1 receptors (Raissi et al., 2013). Plexin-B1 is required in both the presynaptic and postsynaptic neuron for rapid Sema4D-induced synapse formation (McDermott et al., 2018; Adel et al., 2024), suggesting that Sema4D/Plexin-B1 signaling may synchronously coordinate pre- and postsynaptic changes.

In this study, we identify synapse formation events by leveraging the unique ability of Sema4D to drive GABAergic synapse formation with temporal precision combined with two-channel live imaging of fluorescently-labeled GABAergic synaptic proteins. By exploiting Sema4D as a rapid, selective inducer of inhibitory synapse formation, we overcome the inherent asynchrony of synaptogenesis and directly dissect mechanisms that are otherwise difficult to resolve. This work is explicitly focused on the critical time window preceding the emergence of synaptic functionality, thus providing a unique cell biological view of the molecular and structural events that give rise to functional inhibitory synapses. Our results establish a model framework for the cellular processes which underlie inhibitory synapse assembly.

## Results

### Sema4D treatment alters dynamic behavior and promotes stabilization of GAD65-GFP puncta

We first sought to characterize the dynamics of GABAergic presynaptic boutons in response to Sema4D treatment. To begin, we generated primary hippocampal neuronal cultures from P0-1 GAD65-GFP mice, in which a subset of interneurons express a transgenic GAD65-GFP fusion protein (López-Bendito et al., 2004). In these animals axons are identified by diffuse, low-intensity GFP expression along neuronal processes while presynaptic boutons are distinguishable as bright GFP-positive puncta (Fig. 1A, S1, 2A). Previous characterization of this mouse line demonstrated that GAD65-GFP puncta represent genuine presynaptic GABAergic boutons (Wierenga et al., 2008; Poulopoulos et al., 2009; Schuemann et al., 2013; Frias et al., 2019), and we independently confirmed that these GFP-labeled puncta contain GAD65 by immunostaining with a GAD65-specific antibody (Fig. S1).

**Figure 1.**
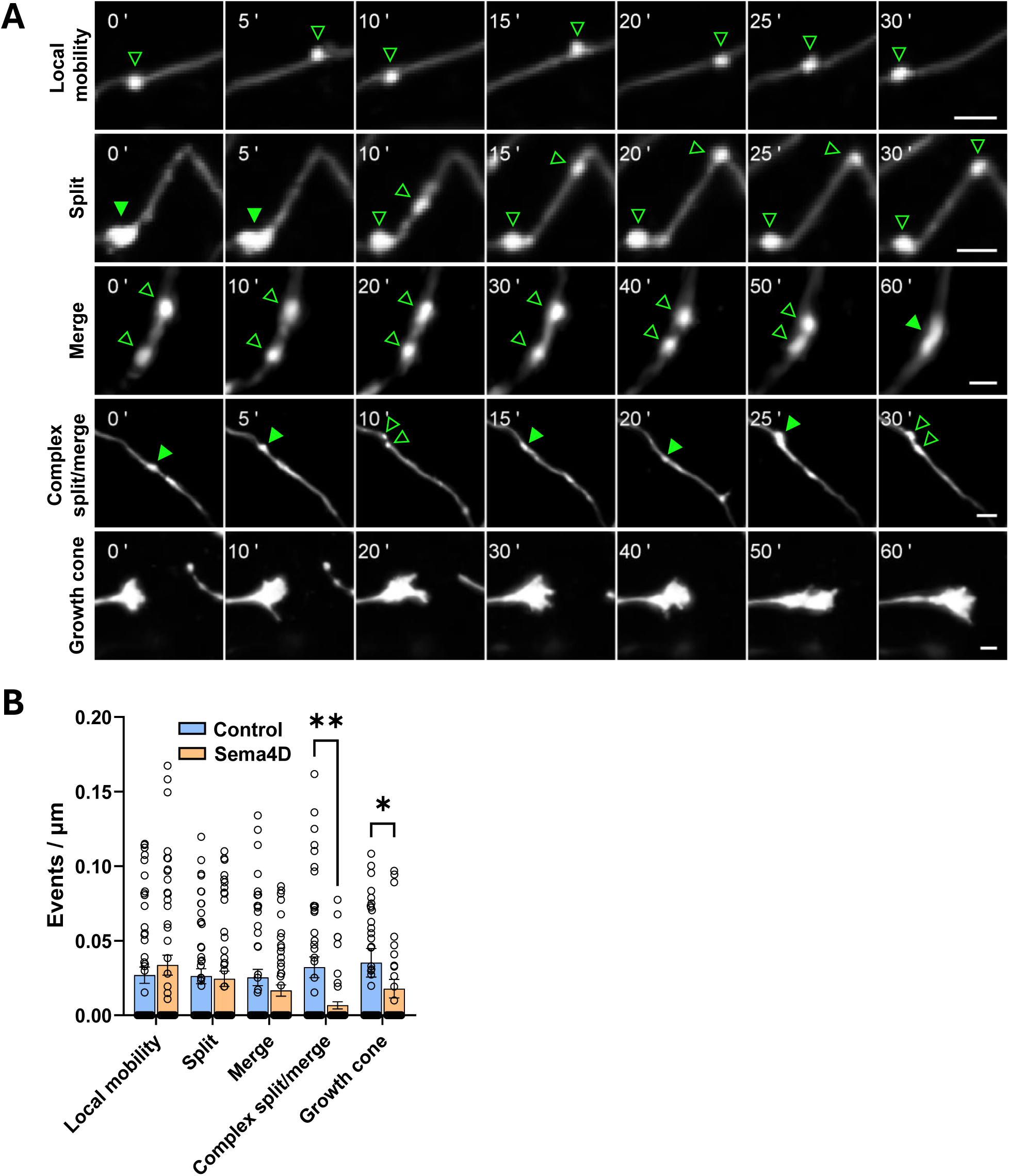
Sema4D affects the frequency of dynamic behaviors of GAD65-GFP puncta in cultured mouse hippocampal neurons. (A) Representative images of stretches of axon from DIV11 GAD65-GFP mouse hippocampal neurons showing dynamic behaviors of interest. (Top to bottom) local mobility (repeated movement of a single cluster along the axon), split, merge, complex split/merge, and growth cone events. Note: time scales differ between events; time points are relative to the start of the montage. Example images include cells from either treatment condition; some montages show different axonal regions from the same neuron. Hollow arrows = single puncta; solid arrows = merged puncta. Scale bars = 2 µm. (B) Frequency of local mobility, split, merge, complex split/merge, and growth cones observed in control vs. Sema4D-treated cultures. Dots correspond to individual axons; n = 50-55 axons per condition from 11 cells (Fc) or 13 cells (Sema4D). *p < 0.05; **p < 0.01; Mann-Whitney U-test.

**Figure 2.**
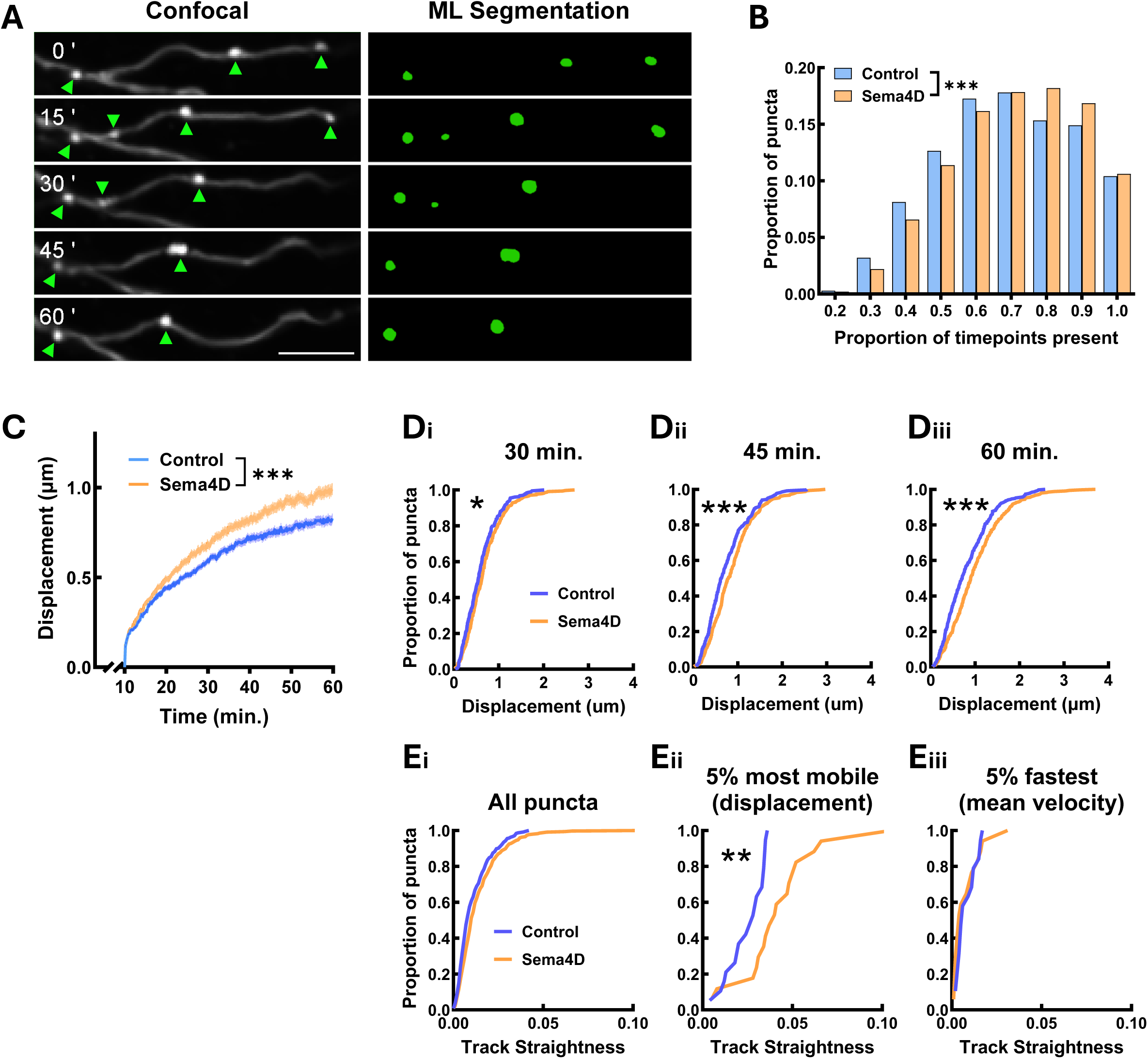
Sema4D treatment increases stability and mobility of presynaptic GAD65-GFP puncta. (A) (Left) Example stretch of axon from cultured hippocampal GAD65-GFP neurons. GAD65-GFP puncta appear as bright spots marked by arrows. (Right) Machine learning (ML) reconstruction of GAD65-GFP puncta in Imaris at the same timepoints. Scale bar = 5 µm. (B) GAD65-GFP puncta stability (proportion of frames in which a puncta was tracked). n = 7427 puncta (Fc), 10213 puncta (Sema4D). The distributions of GAD65-GFP puncta are significantly different between treatment conditions (Chi-square test; χ^2^(8) = 72.66, ***p < 0.001). (C) GAD65-GFP puncta mobility (mean displacement from origin) is increased beginning within 20 min. of Sema4D treatment (binned LME: time × treatment interaction: F(1, 3651) = 11.578, p < 0.001). n = 384 puncta (Fc), 347 puncta (Sema4D). Note: analysis begins at t=10 min. (D) Cumulative frequency histogram of GAD65-GFP puncta displacement at 30, 45, and 60 min. (*p < 0.05, ***p < 0.001; Kolmogorov-Smirnov test). n ≥ 371 puncta per timepoint from 10 cells (Fc), n ≥ 333 puncta per timepoint from 12 cells (Sema4D). (E) Cumulative frequency histogram of GAD65-GFP track straightness. Track straightness was calculated as displacement / total path length. (i) Distribution of track straightness for all puncta (n = 384 puncta (Fc), 347 puncta (Sema4D); p = 0.2594, Kolmogorov-Smirnov test). (ii) Distribution of track straightness for the top 5% most mobile puncta in each condition by displacement (n = 19 puncta (Fc), 17 puncta (Sema4D); **p < 0.01; Kolmogorov-Smirnov test). (iii) Distribution of track straightness for the top 5% of puncta by mean velocity (n = 19 puncta (Fc), 17 puncta (Sema4D); p = 0.8075; Kolmogorov-Smirnov test).

Dissociated GAD65-GFP hippocampal neurons were plated atop an astrocyte feeder layer and cultured for 10 days in vitro (DIV). At DIV10–11, we treated cultures acutely with 2 nM control protein (Fc domain of human IgG_1_ recombinant protein) or 2 nM Sema4D-Fc (recombinant protein containing the soluble extracellular domain of Sema4D fused to the Fc domain of human IgG_1_). Immediately following addition of the recombinant proteins, we acquired images of GAD65-GFP positive neurons using a Nikon AX-R resonant scanning confocal microscope to obtain z-stacks of 5 Nyquist-sampled planes at 10-second intervals for 1 hour.

GAD65-GFP puncta and their associated axons exhibited a range of dynamic behaviors including splitting events, merging events, rapid local protein cluster mobility, complex split/merge events, nascent axonal branching, and active growth cones (Fig. 1A). We manually characterized the behaviors of GAD65-GFP labeled presynaptic protein clusters by identifying and quantifying these events of interest (Fig. 1B; for complete definitions see Table S1). Sema4D treatment significantly decreased the frequency of complex split/merge events, defined as multiple or repeated splitting and merging behavior of one or more GAD65-GFP puncta, compared to control treatment (Fig. 1B). Sema4D treatment also decreased the number of active axonal growth cones observed, consistent with its previously demonstrated role in growth cone collapse (Oinuma et al., 2004; Ito et al., 2006), thus confirming that Sema4D-Fc protein is active in our cultures. By contrast the rates of single split, single merge, and locally restricted mobility events (mobile puncta moving within a small radius without splitting or merging) of GAD65-GFP puncta, as well as the rate of nascent axonal branch formation, were unaffected. This suggests that Sema4D signaling may have a role in stabilizing a specific subset of immature, mobile presynaptic protein assemblies that undergo repeated split/merge events.

To track the mobility of GAD65-GFP puncta quantitatively, we identified puncta at each timepoint using a custom-trained machine learning segmentation model in the Imaris for Tracking suite (Oxford Instruments) and empirically determined that this rigorous approach produced the highest fidelity puncta identification over time (Fig. 2A). We then performed automated particle tracking using Imaris and a custom analysis suite in MATLAB. We quantified GAD65-GFP puncta stability by calculating the percentage of imaging frames in which each individual puncta was present, similar to prior studies of GAD65-GFP boutons in organotypic slice (Frias et al., 2019). Sema4D treatment shifted the distribution of puncta duration rightward (Fig. 2B), indicating that GAD65-GFP puncta were present at a greater proportion of timepoints. Thus, Sema4D treatment appears to stabilize existing GAD65-GFP puncta.

### GAD65-GFP puncta show altered mobility in response to Sema4D treatment

To characterize the mobility of individual GAD65-GFP puncta in response to Sema4D we focused on GAD65-GFP puncta that were consistently tracked between t = 10 min. and t = 60 min. of the imaging session. Beginning the analysis at t = 10 min. was necessary due to an initial, transient rise in mean displacement distance for all GAD65-GFP puncta within this time window in both treatment conditions, presumably due to addition of the protein to the culture media or imaging-induced cell motility. We first characterized overall GAD65-GFP puncta mobility by tracking displacement from origin of every identified GAD65-GFP puncta in the field of view starting at t = 10 min. We found that Sema4D treatment led to an overall increase in mean GAD65-GFP puncta mobility beginning about 20 minutes after addition of Sema4D protein, as shown by a significant increase in mean displacement in Sema4D-treated cultures compared to control from approximately 20–60 min. (Fig. 2C). We next compared the overall distributions of GAD65-GFP puncta mobility across the population at 30, 45, and 60 min. after Sema4D or control treatment. Across both treatment conditions, the majority (approximately 90%) of GAD65-GFP puncta were relatively immobile, displacing less than 2 µm from their initial position, consistent with the presence of stable protein clusters localized to presynaptic boutons (Fig. 2D). We observed that overall mean displacement was significantly increased in Sema4D-treated cultures at these timepoints (Fig. 2D). However, the shape of the cumulative distribution was similar between treatment conditions, suggesting that Sema4D induces a modest, population-wide shift toward greater puncta mobility rather than actuating a distinct subpopulation of highly mobile puncta.

Despite the similarity in overall population distribution, the most mobile GAD65-GFP puncta moved a greater distance in Sema4D-treated neurons compared to control across all timepoints (Fig. 2D). We therefore asked whether Sema4D-dependent changes in GAD65-GFP puncta mobility were partly due to increased directionality (i.e. track straightness) of mobile GAD65-GFP puncta (Fig. S2A), as mean and maximum GAD65-GFP puncta velocity did not differ between conditions (Fig. S2B, C). Analysis of track straightness revealed that while Sema4D treatment did not affect the average track straightness of the entire population of puncta (Fig. 2Ei), the most mobile subset of GAD65-GFP puncta (top 5% by displacement within each condition) followed significantly straighter tracks in the Sema4D-treated cultures (Fig. 2Eii). By contrast, track straightness of the top 5% of GAD65-GFP puncta by mean velocity did not differ between treatment conditions (Fig. 2Eiii). Interestingly, the autoregressive parameter AR(1), which represents persistence in track directionality, was increased in the top 5% of GAD65-GFP puncta by mean velocity in Sema4D-treated cultures, suggesting that faster-moving GAD65-GFP puncta moved more freely under Sema4D treatment (Fig. S2D). Overall, these data suggest that Sema4D treatment drives more directed movement among a mobile subset of GAD65-GFP puncta, and that increased GAD65 puncta displacement may in part be driven by more directional movement of a subset of mobile presynaptic protein clusters.

Stabilization of presynaptic protein clusters at new synaptic sites may be mediated by altered mobility of existing protein clusters as well as maturation of unstable, newly-formed clusters. A prior live imaging study with a 10-minute temporal resolution reported that Sema4D stabilizes a subset of non-persistent boutons which were intermittently present before treatment (Frias et al., 2019). Thus, while our current analysis was confined to GAD65-GFP boutons that were tracked stably across the entire 60-minute imaging session, we also considered the possibility that Sema4D may act on the newly-formed GAD65-GFP puncta that appeared after the onset of imaging. To test whether Sema4D affects the mobility of this subset, we focused on GAD65-GFP puncta that emerged between t = 3 and t = 20 min. after imaging onset and remained stably present thereafter. While these newly-tracked puncta were generally more mobile than existing puncta (Fig. S2E), we did not observe a Sema4D-dependent effect on mean displacement of newly-tracked puncta at any timepoint (Fig. S2F).

Overall, the results from this imaging experiment indicate that a subset of GAD65-GFP protein clusters are mobile and display dynamic behaviors during the 1 hr. imaging session. Sema4D-dependent changes to presynaptic GAD65-GFP puncta mobility are time-dependent, with an overall increase in mean puncta mobility beginning about 20 min. after Sema4D addition. Increased mobility is accompanied by increased puncta stability and a decrease in the frequency of complex split/merge events, suggesting that Sema4D promotes both increased exploratory dynamics and maturation of presynaptic sites.

### Sema4D does not affect overall mobility of postsynaptic gephyrin

We next asked whether Sema4D similarly affects the mobility of postsynaptic proteins associated with GABAergic synapses. We generated primary hippocampal cultures from P0–1 GAD65-GFP mice as described above. At DIV2 these cultures were infected with an AAV9 virus expressing HaloTag-Gephyrin under the control of the neuronal hSyn promoter (Halo-Gephyrin), allowing for visualization of postsynaptic scaffold assemblies (Fig. 3A; Fig. S3A). We confirmed that virally expressed Halo-Gephyrin localizes to synapses as revealed by co-immunostaining with a GAD65-specific antibody (Fig. S3A), and while gephyrin puncta density is marginally increased in these cultures, baseline GABAergic synapse density was unaffected (Fig. S4). At DIV10–11 we labeled cultures with cell-permeable Janelia Fluor 646 (JF646) HaloTag ligand to visualize Halo-Gephyrin, then treated with 2 nM Sema4D or control protein and acquired images at 15s intervals for up to 1h using a resonant scanning confocal. As before, we utilized a custom-trained machine learning segmentation network, automated particle tracking in Imaris, and custom MATLAB-based analysis software to analyze the mobility of Halo-Gephyrin puncta (Fig. 3A). In contrast to GAD65-GFP puncta, we observed no effect of Sema4D treatment on Halo-Gephyrin puncta stability over time (Fig. 3B). Similar to GAD65-GFP, 90–95% of Halo-Gephyrin puncta were relatively immobile, displacing less than 2 µm from their starting location (Fig. 3D). However, we did not observe a Sema4D-dependent effect on the distribution of Halo-Gephyrin puncta mobility at any timepoint (Fig. 3D), nor did we observe any appreciable Sema4D-dependent changes to Halo-Gephyrin puncta velocity (Fig. S3B, C) or to indicators of directed motion such as gephyrin track straightness or autoregressive parameter (Fig. 3E, S3D). Thus, we concluded that Sema4D does not affect the overall mobility of gephyrin scaffolds within our imaging window. Taken together these data suggest that Sema4D promotes GABAergic synapse formation primarily by altering the behavior of the presynaptic terminal.

**Figure 3.**
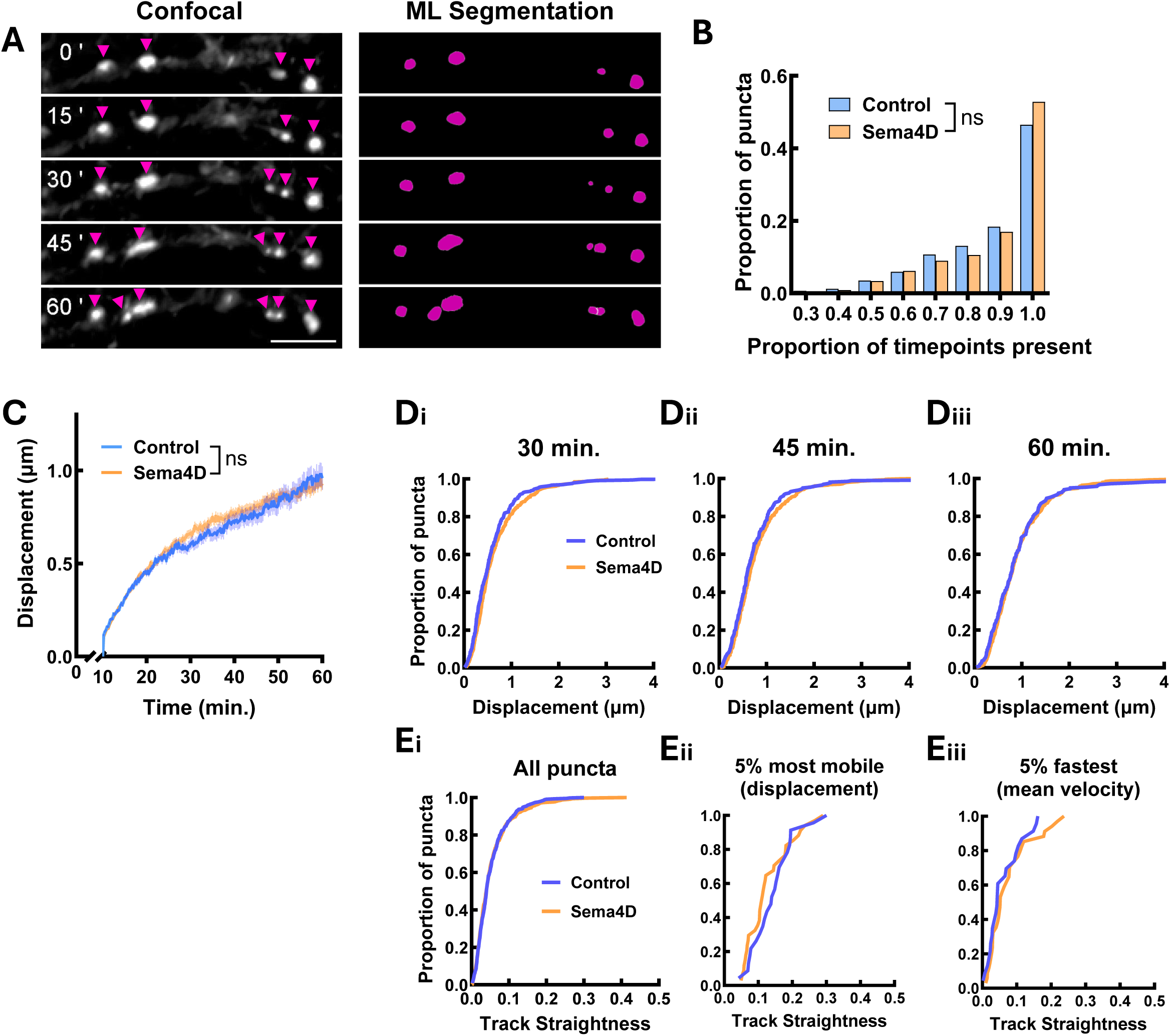
Sema4D treatment does not affect Halo-Gephyrin mobility at the population level. (A) Representative stretch of dendrite showing Halo-Gephyrin expression along dendrites in cultures from GAD65-GFP neurons. Halo-Gephyrin puncta appear as bright spots marked by arrows. (Right) Machine learning (ML) reconstruction of Halo-Gephyrin puncta in Imaris at the same timepoints. Scale bar = 5 µm. (B) Halo-Gephyrin puncta stability (proportion of frames in which a puncta was tracked). n = 742 puncta (Fc), 1026 puncta (Sema4D). The distributions of Halo-Gephyrin puncta do not significantly differ between treatment conditions (Chi-square test; χ^2^ (7) = 10.63, p = 0.1557). (C) Halo-Gephyrin puncta mobility (mean displacement from origin) is not affected by Sema4D treatment (binned LME: time × treatment interaction: F(1, 7279) = 1.0464, p = 0.2954). Note: analysis begins at t=10 min. (D) Cumulative frequency histogram of Halo-Gephyrin puncta displacement at 30, 45 and 60 min. There was no effect of Sema4D treatment on the overall distribution of Halo-Gephyrin displacement at 30 min. (p = 0.0529; Kolmogorov-Smirnov test), at 45 min. (p = 0.3440), or at 60 min. (p = 0.7637). n ≥ 220 puncta per timepoint from 8 cells (Fc), n ≥ 393 puncta per timepoint from 8 cells (Sema4D). (E) Cumulative frequency histogram of Halo-Gephyrin track straightness. (i) Distribution of track straightness for all puncta (n = 384 puncta (Fc), 347 puncta (Sema4D); p = 0.8307, Kolmogorov-Smirnov test). (ii) Distribution of track straightness for the top 5% most mobile puncta in each condition by displacement (n = 23 puncta (Fc), 34 puncta (Sema4D); p = 0.3308; Kolmogorov-Smirnov test). (iii) Distribution of track straightness for the top 5% of puncta by mean velocity (n = 23 puncta (Fc), 34 puncta (Sema4D); p = 0.6355; Kolmogorov-Smirnov test).

### Sema4D treatment drives gephyrin localization to postsynaptic sites adjacent to GAD65-GFP labeled boutons

Our previous studies using immunostaining in fixed cells of proteins localized to GABAergic synapses demonstrated that Sema4D promotes increased density of GAD65 puncta colocalized with GABA_A_Rγ2 and of synapsin puncta colocalized with gephyrin (Kuzirian et al., 2013). Here we sought to determine the time course of colocalization between pre- and postsynaptic markers of GABAergic synapses that occurs in response to Sema4D. We first asked whether Sema4D treatment promotes increased localization of Halo-Gephyrin to sites marked by GAD65-GFP puncta. For this analysis we focused primarily on stably tracked GAD65-GFP boutons that were present throughout the 1 hr. imaging session. Because GAD65 and gephyrin show largely overlapping point-spread functions at standard confocal resolution, we used this overlapping signal as an indicator of colocalization between Halo-Gephyrin postsynaptic puncta and GAD65-GFP labeled presynaptic boutons (Fig. 4A). We observed an increase in the mean fluorescence intensity of the Halo-Gephyrin channel within the area defined by GAD65-GFP puncta (hereafter referred to as “colocalized gephyrin fluorescence”) across time. At the population level, we found that Sema4D treatment significantly increased mean colocalized gephyrin fluorescence beginning at about 45 min. (Fig. 4Bi), consistent with our previous results in fixed cells.

**Figure 4.**
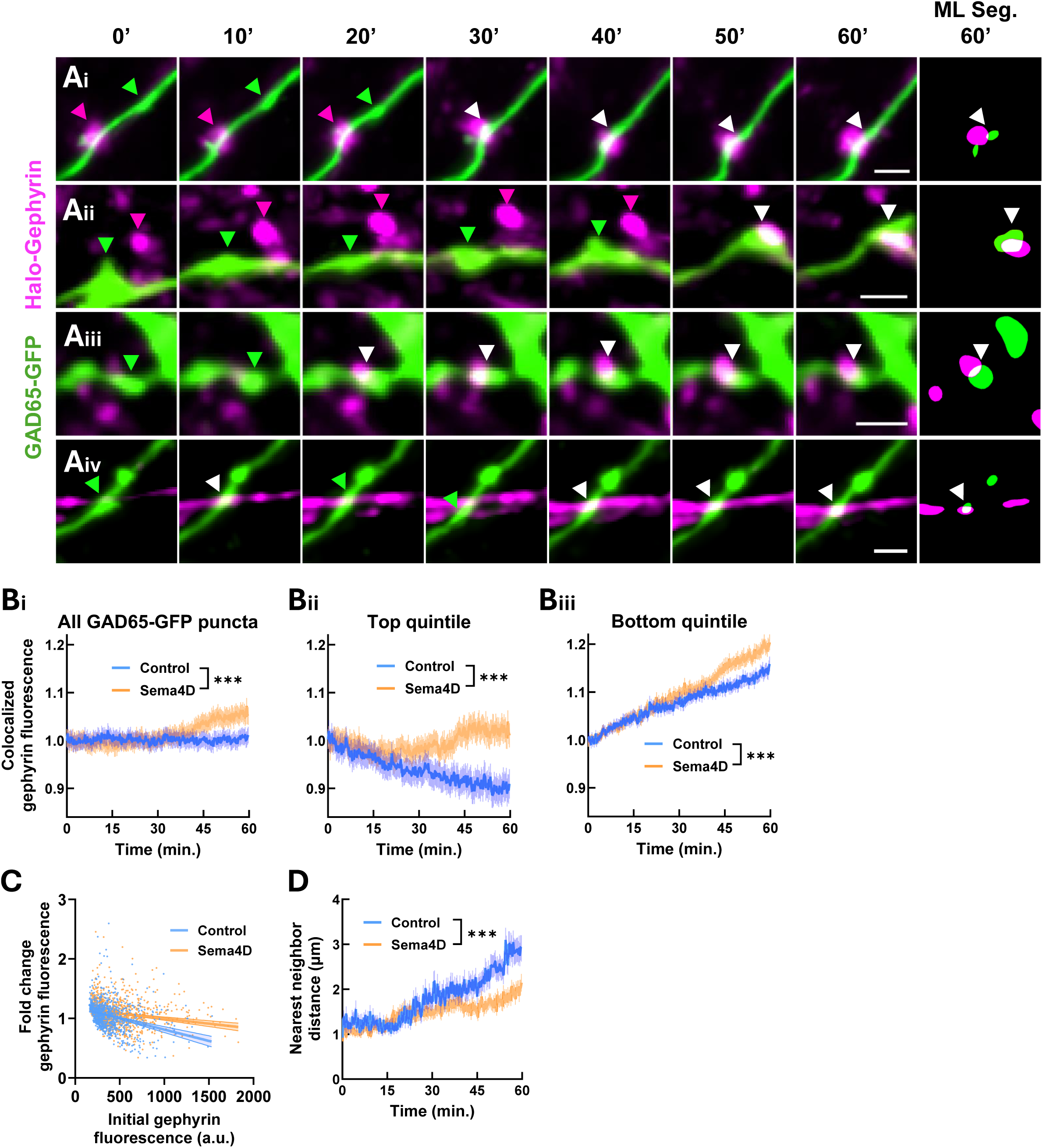
Sema4D promotes increased gephyrin colocalization with existing GAD65-GFP puncta. (A) Montages showing putative new synapse formation events. (i, ii) Existing mobile GAD65-GFP puncta (green arrows) localize to existing Halo-Gephyrin puncta (magenta arrows) and remain colocalized (white arrows). (iii, iv) A newly-tracked Halo-Gephyrin puncta emerges and colocalizes with an existing GAD65-GFP puncta (green arrows). White arrows = colocalized puncta. Scale bars = 2 µm. (B) (i) Gephyrin fluorescence colocalized with GAD65-GFP puncta is increased in Sema4D condition compared to control (binned LME: time × treatment interaction: F(1, 24140) = 13.656, ***p < 0.001). n ≥ 765 puncta per timepoint (control), 1000 puncta (Sema4D). (ii) Sema4D prevents the loss of gephyrin from GAD65-GFP boutons in the top quintile of baseline gephyrin signal (interaction : F(1, 4820) = 17.75, ***p < 0.001). n ≥ 151 puncta per timepoint (control), 201 puncta (Sema4D). (iii) Sema4D treatment increases recruitment of gephyrin to boutons in the bottom quintile of baseline gephyrin signal compared to control treatment (interaction : F(1, 4820) = 17.637, ***p < 0.001). n ≥ 150 puncta per timepoint (control), 197 puncta (Sema4D). Error bars = SEM. Data are normalized within treatment condition to the mean of the first 3 minutes. (C) There is a negative correlation between baseline (t=0 min.) gephyrin fluorescence colocalized with GAD65-GFP puncta and change in gephyrin fluorescence from t=0 min. to 60 min; Sema4D treatment reduces the strength of this correlation. Control: slope CI = (−4.568×10^-4^, -3.039×10^-4^); Sema4D: slope CI = (−2.150×10^-4^, -1.241×10^−4^). n = 866 puncta (control), 1146 puncta (Sema4D). Error bars = 95% CI of linear regression fit. (D) Sema4D treatment decreases the mean nearest neighbor distance of GAD65-GFP puncta to the nearest gephyrin puncta (for GAD65-GFP puncta with initial nearest neighbor distance ≤ 2 µm) (binned LME: time × treatment interaction : F(1,4976) = 22.334, ***p < 0.001). n ≥ 123 puncta per timepoint (control), 242 puncta (Sema4D). Error bars = SEM.

To assess whether Sema4D-dependent recruitment of gephyrin to GAD65-GFP puncta varied as a function of the initial degree of postsynaptic association, we analyzed subsets of GAD65-GFP puncta grouped by their baseline levels of colocalized gephyrin fluorescence. The top quintile, characterized by greater colocalized gephyrin fluorescence at t = 0 min., showed a gradual decrease in colocalized gephyrin fluorescence over time in the control condition; Sema4D treatment prevented the loss of gephyrin fluorescence from this subset (Fig. 4Bii). By contrast, the bottom quintile of GAD65-GFP puncta, characterized by little-to-no colocalized gephyrin fluorescence initially, showed gradually increasing colocalized gephyrin fluorescence over time in control and Sema4D-treated neurons. Sema4D treatment led to further increased colocalized gephyrin fluorescence beyond the level seen in control neurons, suggesting additional recruitment of gephyrin to this subset of GAD65-GFP puncta in response to Sema4D treatment (Fig. 4B, iii).

Across the entire dataset, we found a negative correlation between baseline gephyrin fluorescence and the change in gephyrin fluorescence from t = 0 to t = 60 min.; as expected, Sema4D treatment significantly reduced the strength of this correlation (Fig. 4C). Overall, these data suggest that Sema4D acts to 1) stabilize and prevent the loss of gephyrin from sites of colocalization with GAD65 and 2) promote increased colocalization of gephyrin at GAD65-positive sites lacking gephyrin.

We hypothesized that increased colocalization between GAD65-GFP and Halo-Gephyrin puncta could occur via migration of existing GAD65-GFP or Halo-Gephyrin puncta to new sites (Fig. 4Ai,ii) and/or via accumulation of Halo-Gephyrin at sites where GAD65-GFP was already present (Fig. 4Aiii,iv). To examine these possibilities, we measured the mean nearest neighbor distance of GAD65-GFP puncta, defined as the minimum distance between a GAD65-GFP puncta and the nearest Halo-Gephyrin puncta at any given time. Because over 90% of persistently-tracked GAD65-GFP and Halo-Gephyrin puncta displaced 2 µm or less (Figs. 2, 3), we focused on GAD65-GFP puncta that were initially within 2 µm of a gephyrin puncta. Sema4D treatment decreased the mean nearest neighbor distance of GAD65-GFP boutons beginning at about 45 min. compared to control treatment (Fig. 4D). We hypothesized that increased proximity between GAD65 and gephyrin resulted from new colocalization events in which previously non-colocalized GAD65-GFP and Halo-Gephyrin puncta became colocalized with each other during the imaging session.

We next used our two-channel particle tracking data to identify new colocalization events between GAD65 and gephyrin. These events were considered genuine new colocalization events only if they produced stable new colocalization between a pair of GAD65-GFP and Halo-Gephyrin puncta that persisted for at least 10 minutes. New colocalization events fell into two distinct categories: colocalization events between existing GAD65-GFP puncta and newly-emerging Halo-Gephyrin puncta (“new gephyrin”) and colocalization events between existing GAD65-GFP puncta and stable, pre-existing gephyrin puncta (“existing gephyrin”). We found that in both control and Sema4D-treated neurons, colocalization events with new gephyrin puncta comprised about 75% of the total new colocalization events (Fig. 5A). Sema4D treatment significantly increased the frequency of colocalization events between GAD65-GFP and new gephyrin puncta but not between GAD65-GFP and existing gephyrin puncta (Fig. 5B). These data suggest that one way in which colocalized gephyrin fluorescence is increased in response to Sema4D treatment is via new colocalization events between new gephyrin puncta and existing GAD65-labeled sites.

**Figure 5.**
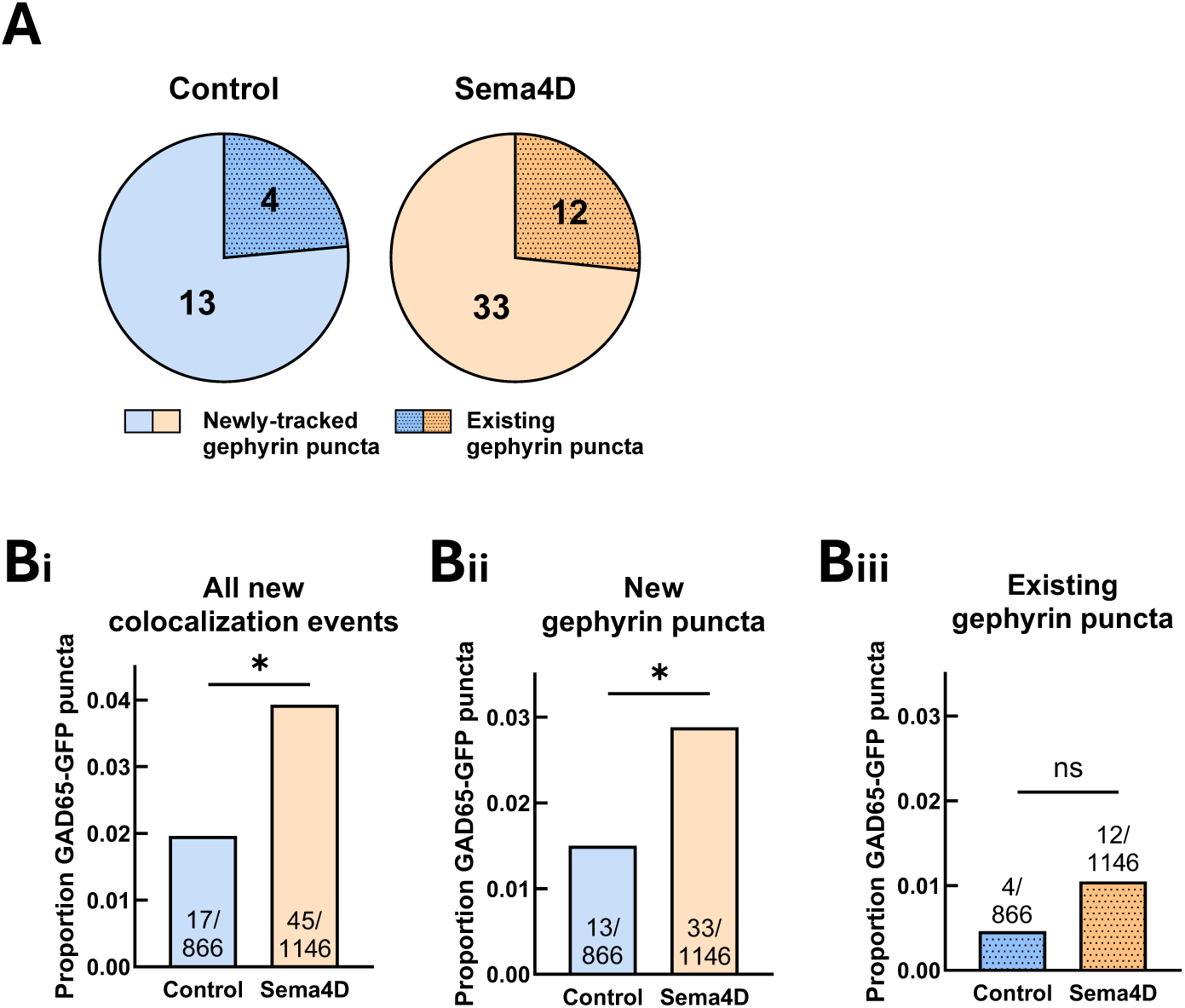
Sema4D-dependent GAD65/gephyrin colocalization occurs via colocalization events between existing GAD65 clusters and new gephyrin clusters. (A) Number of GAD65-GFP puncta that colocalized with an existing Halo-Gephyrin puncta vs. a newly-tracked Halo-Gephyrin puncta. (B) (i) Overall frequency of all new colocalization events. Sema4D treatment significantly increased the frequency of new colocalization events compared to control (*p < 0.05). (ii) Sema4D treatment significantly increased the proportion of GAD65-GFP puncta that colocalized with a newly-tracked Halo-Gephyrin puncta (*p < 0.05, Fisher’s exact test). (iii) There was no significant difference in the frequency of colocalization events with an existing Halo-Gephyrin puncta in Sema4D vs. control-treated cultures (p = 0.2048).

### GAD65-GFP puncta that undergo new colocalization events show distinctive profiles of mobility and proximity to gephyrin puncta

To better understand the spatiotemporal dynamics of GAD65 protein clusters that undergo new colocalization events, we used principal component analysis (PCA) to determine which particle tracking parameters most strongly characterize the variability in individual GAD65-GFP puncta (Table S2). We reasoned that GAD65-GFP puncta that undergo new colocalization events could comprise a distinctly identifiable subset of puncta in principal component space based on a distinctive profile of mobility and proximity to gephyrin puncta. We therefore compared their distribution along PC1 and PC2, the principal axes capturing the greatest variance in puncta features, to that of the overall population. PC1 loadings were highest for measures of GAD65-GFP puncta mobility, including mean velocity and acceleration, while PC2 loadings were highest for measures of GAD65-GFP proximity to gephyrin puncta, including initial and mean nearest neighbor distance to gephyrin. We found that GAD65-GFP puncta that undergo colocalization events tend to be relatively immobile and are located relatively near to existing Halo-Gephyrin puncta (Fig. S5A). Surprisingly, GAD65-GFP puncta that colocalized with new gephyrin puncta showed no difference in average velocity compared to other GAD65-GFP puncta in either control or Sema4D-treated cultures, suggesting that additional aspects of mobility are required to predict new colocalization events (Fig. S5B). In each treatment condition, mean initial nearest neighbor distance to gephyrin was significantly lower for GAD65 puncta with a new colocalization event, suggesting that even though these puncta colocalized with gephyrin puncta that emerged later during the imaging session, local proximity to other existing gephyrin puncta at the onset of imaging was strongly predictive of new colocalization events (Fig. S5C). This suggests that GAD65 puncta undergoing colocalization events are localized to “hot spot” regions of synapse assembly where more gephyrin is present to be recruited from nearby sites to form new synapses.

We performed the same analysis for new colocalization events in which GAD65-GFP puncta colocalized with an existing Halo-Gephyrin puncta. These events were relatively less common, only comprising about 25% of new colocalization events (Fig. 5A); due to the smaller sample size, we could not separately analyze control and Sema4D treatment conditions for this subset of events. Similar to GAD65-GFP puncta that colocalized with new gephyrin puncta, the GAD65-GFP puncta that colocalized with existing gephyrin puncta displayed lower average mobility and greater initial proximity to gephyrin compared to other GAD65-GFP puncta (Fig. S5D). These GAD65-GFP puncta had significantly lower mean velocity compared to puncta without a colocalization event (Fig. S5E), and initial nearest neighbor distance to existing gephyrin was zero in every case in which we observed a new colocalization event (Fig. S5F), suggesting that these GAD65 puncta were initially colocalized with existing gephyrin, temporarily lost contact, and then colocalized again.

Taken together, these results suggest that features of GAD65-GFP puncta such as mobility and proximity to postsynaptic sites marked by gephyrin are partially predictive of whether stable new colocalization will occur. Sema4D treatment significantly increased the overall probability of new colocalization events, primarily by enhancing the formation of stable associations with newly emerging gephyrin puncta. Overall, these analyses indicate that GAD65 boutons that form new, stable colocalization events are defined primarily by their initial proximity to gephyrin-rich regions, which strongly predicts whether they will recruit new postsynaptic scaffold.

### Sema4D treatment mobilizes GAD65-GFP puncta prior to colocalization with Halo-Gephyrin

Given that Sema4D increases the overall mobility of GAD65-GFP but not Halo-Gephyrin puncta, and that Sema4D increases the probability of new colocalization events between GAD65 and gephyrin, we hypothesized that presynaptic protein clusters define the sites of new GABAergic synapses. To test this directly we performed an event-level analysis of paths followed by GAD65-GFP or Halo-Gephyrin puncta before and after new colocalization events (Fig. 6A, E). For this analysis, all new colocalization events (including those involving new and existing gephyrin puncta) were combined for each treatment condition. GAD65-GFP puncta that colocalized with Halo-Gephyrin puncta displaced significantly farther from their origin prior to colocalization in Sema4D-treated cultures compared to control (Fig. 6B). However, when we examined the mean distance of GAD65-GFP puncta from the site of colocalization specifically in the 10-minute window prior to colocalization events, there was no difference between control and Sema4D-treated cultures (Fig. 6C), suggesting that Sema4D-dependent changes to GAD65 mobility occur in an earlier window relative to when GAD65 puncta become colocalized with gephyrin.

**Figure 6.**
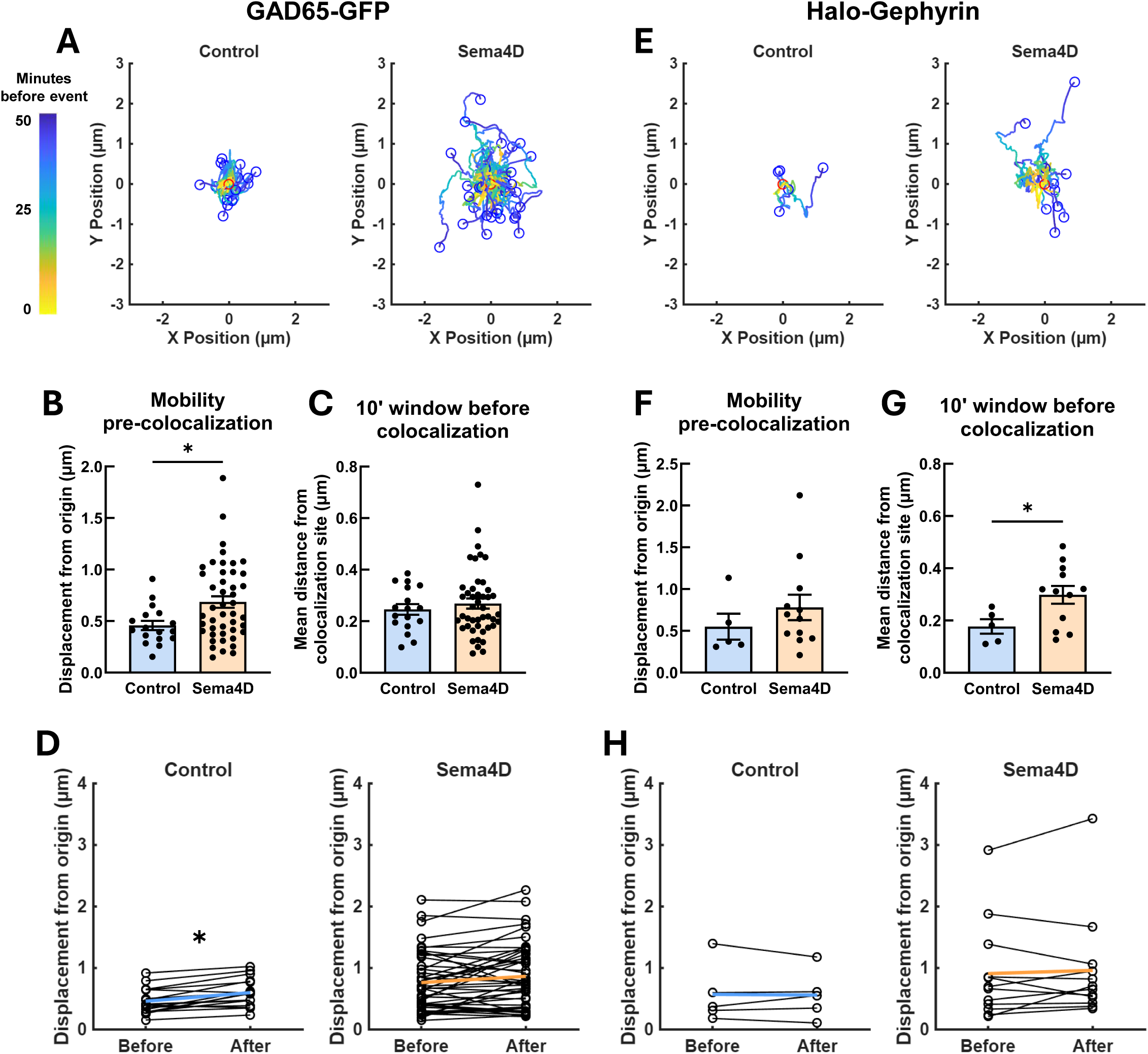
Sema4D-dependent GAD65/gephyrin colocalization events are driven by early GAD65-GFP displacement followed by late Halo-Gephyrin mobility. (A) Tracks followed by GAD65-GFP puncta prior to new colocalization events with gephyrin in control (left) or Sema4D (right)-treated cultures. Blue dots represent relative start locations; red dots (at 0,0) represent normalized location of new colocalization event. (B) Total displacement of GAD65-GFP puncta undergoing a new colocalization event across entire imaging session (*p < 0.05, Mann-Whitney U-test). n = 17 events (control), 45 events (Sema4D). Error bars = SEM. (C) Mean distance of GAD65-GFP puncta from new colocalization sites in the 10-minute window preceding new colocalization events. Sema4D treatment did not affect mean GAD65-GFP distance during this time window (p = 0.8513, Mann-Whitney U-test). n = 17 events (control), 45 events (Sema4D). Error bars = SEM. (D) Change in mean displacement from origin of individual GAD65-GFP puncta before vs. after colocalization. GAD65-GFP puncta are significantly further from origin after colocalization events in control neurons (*p < 0.05, Wilcoxon matched-pairs signed rank test) but not in Sema4D-treated neurons (p = 0.0973) . n = 17 events (control), 45 events (Sema4D). Error bars = SEM. (E) Tracks followed by Halo-Gephyrin puncta prior to new colocalization events with GAD65-GFP in control (left) or Sema4D (right) treated cultures. Blue dots represent relative start locations; red dots (0,0) represent normalized location of new colocalization event. (F) Total displacement of Halo-Gephyrin puncta undergoing a new colocalization event across entire imaging session. Sema4D treatment does not affect displacement of Halo-Gephyrin puncta undergoing colocalization events (p = 0.2343, Mann-Whitney U-test). n = 5 events (control), 12 events (Sema4D). Error bars = SEM. (G) In Sema4D-treated cultures Halo-Gephyrin puncta are on average significantly farther from new colocalization sites during the 10-minute window preceding new colocalization events compared to control cultures (*p < 0.05, Mann-Whitney U-test). n = 5 events (control), 12 events (Sema4D). Error bars = SEM. (H) Mean displacement from origin of individual Halo-Gephyrin puncta in the 10-minute window before vs. after colocalization. Halo-Gephyrin does not travel significantly further from origin following colocalization events in either control (p = 0.999, Wilcoxon matched-pairs signed rank test) or Sema4D-treated neurons (p = 0.677). n = 5 events (control), 12 events (Sema4D). Error bars = SEM.

To determine whether GAD65-GFP puncta are moving towards pre-determined locations to establish new colocalization sites, we analyzed the change in GAD65-GFP displacement from origin during the 10-minute window before and the 10-minute window after colocalization. We reasoned that if a new colocalization event was driven by a GAD65-GFP puncta moving to a pre-determined location along the neurite to colocalize with a gephyrin puncta (or vice versa), GAD65-GFP puncta would show greater average displacement after colocalization vs. before colocalization, indicating the GAD65-GFP puncta had moved farther from its origin towards the pre-determined location. GAD65-GFP puncta displacement was slightly (albeit significantly) increased after colocalization in control neurons, but not in Sema4D-treated neurons, confirming that the Sema4D-dependent increase to GAD65-GFP puncta mobility occurs during an earlier window (Fig. 6D). In addition, because Sema4D treatment did not significantly change GAD65-GFP displacement after colocalization vs. before, we conclude that Sema4D treatment does not direct GAD65-GFP towards specific pre-determined locations along the axon. Notably, in the Sema4D-treated condition, several GAD65-GFP puncta moved more than 1 µm prior to a new colocalization event, a phenomenon which was not observed in control cultures (Fig. 6A, D).

We next performed the same analysis for Halo-Gephyrin puncta that became stably colocalized with a GAD65-GFP puncta. In contrast to GAD65-GFP puncta, Sema4D treatment had no effect on Halo-Gephyrin puncta displacement prior to colocalization (Fig. 6F), although Halo-Gephyrin puncta were on average farther from the colocalization site in the 10-minute window prior to colocalization in Sema4D-treated cultures compared to control (Fig. 6G). This suggests that gephyrin puncta mobility is influenced by Sema4D signaling in the immediate window prior to colocalization, but not before, possibly because the presence of a nearby presynaptic bouton is required for the gephyrin puncta to mobilize. Halo-Gephyrin puncta also did not show greater displacement from origin after colocalizing with GAD65-GFP vs. before colocalizing in either treatment condition, suggesting that gephyrin scaffold mobility is not specifically directed towards pre-determined locations along dendrites (Fig. 6H). Taken together, these results suggest that GAD65-GFP mobility occurs first in the sequence of events underlying Sema4D-dependent synapse formation, followed by later gephyrin recruitment, and that mobile GAD65-GFP puncta likely encounter gephyrin assemblies stochastically.

### Recruitment of GABA_A_Rγ2 to gephyrin scaffolds is increased in response to Sema4D treatment

Recruitment and stabilization of GABA_A_Rs at postsynaptic gephyrin scaffolds is an essential step in GABAergic synapse maturation, and gephyrin has been shown to regulate GABA_A_R clustering and surface expression (Mukherjee et al., 2011; Petrini et al., 2014). Previous work from our lab demonstrated that the density of GABA_A_Rγ2-positive synapses is increased in response to Sema4D treatment and that newly-formed GABAergic synapses are functional within 2 hours (Kuzirian et al., 2013), suggesting that functional GABA_A_Rs are localized to newly-formed synapses. However, gephyrin–GABA_A_R colocalization has not been assessed directly in response to Sema4D treatment, and the spatiotemporal dynamics of their association during synapse formation are poorly understood. To this end, we co-transfected plasmids expressing GFP-Gephyrin and HaloTag-GABA_A_Rγ2 subunit (Halo-γ2) into cultured wild-type E18 rat neurons at DIV 4 and performed live imaging at DIV10–11 as before using cell-permeable JF646 HaloTag ligand to label the Halo-γ2 protein (Figs. S6A, 7A). As with the virally expressed Halo-Gephyrin construct, we observed discrete clusters of GFP-Gephyrin, whereas Halo-γ2 expression was significantly more variable. In line with prior observations (Christie et al., 2006), a subset of GFP-Gephyrin/Halo-γ2 co-transfected neurons showed both dispersed and punctate Halo-γ2 localization along dendrites (Fig. S6A), with the remaining co-transfected neurons showing only weakly dispersed Halo-γ2 signal in dendrites, perhaps representing less mature or more weakly transfected neurons. GFP-Gephyrin and Halo-γ2 puncta generally colocalized, and fixed immunostaining experiments showed that overexpression of these markers did not affect baseline GABAergic synapse density or interfere with synaptic localization of Halo-γ2 puncta (Fig. S7, S8). Thus, we proceeded with live imaging analysis for co-transfected cells with clear punctate Halo-γ2 fluorescence.

We first analyzed the mean fluorescence intensity of Halo-γ2 colocalized with GFP-Gephyrin as a measure of receptor density at postsynaptic scaffolds (Fig. 7B), similar to our previous analysis of GAD65 and gephyrin colocalization (Fig. 4). We found that the mean colocalized Halo-γ2 fluorescence rapidly increased in response to Sema4D treatment within about 10 minutes of application. By 1 hour, the mean increase in colocalized Halo-γ2 fluorescence puncta was approximately 10–15% above baseline in Sema4D treated cultures (Fig. 7Bi). Sema4D treatment did not affect the mean intensity of individual Halo-γ2 puncta or the total dendritic intensity in the Halo-γ2 channel (Fig. S6B, C); thus, the increase in colocalized Halo-γ2 fluorescence is likely due to increased recruitment of Halo-γ2 to gephyrin-positive postsynaptic scaffolds rather than increased GABA_A_R expression.

**Figure 7.**
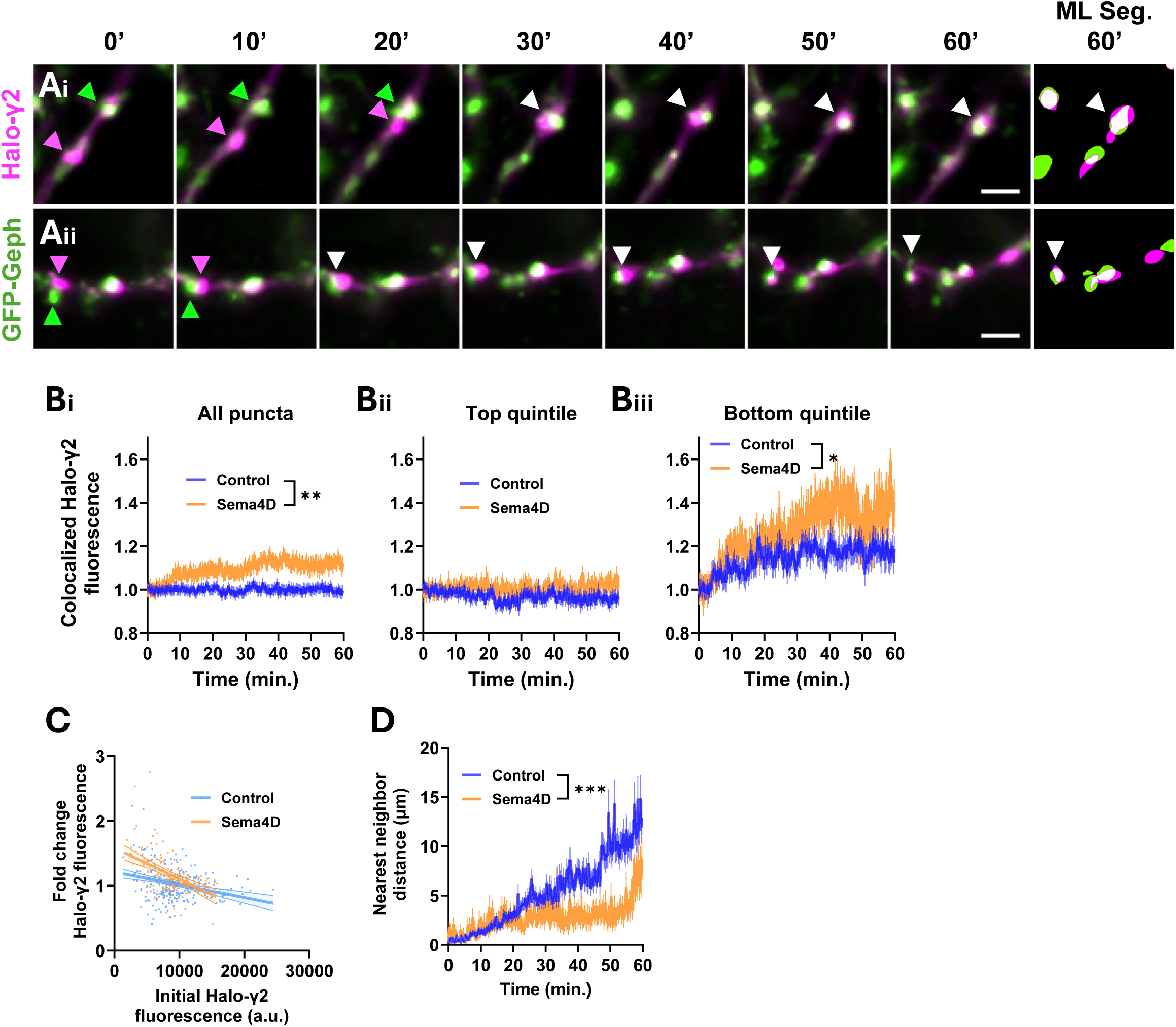
Sema4D promotes increased localization of GABA_A_Rγ2 to gephyrin-labeled postsynaptic specializations. (A) Representative images showing colocalization between GFP-Gephyrin (green arrows) and Halo-γ2 (magenta arrows). (i, ii) Examples showing a Halo-γ2 puncta moving and colocalizing with a GFP-Gephyrin puncta (white arrows = colocalization). Images represent different regions from the same neuron. Scale bars = 2 µm. (B) (i) Sema4D treatment increases mean Halo-γ2 intensity colocalized with GFP-Gephyrin puncta compared to control treatment (binned LME: time × treatment interaction: F(1, 4964) = 8.5686, **p < 0.01). n ≥ 254 puncta per timepoint (control), 130 puncta (Sema4D). (ii) Sema4D treatment does not affect mean Halo-γ2 intensity colocalized with GFP-Gephyrin puncta in the top quintile of baseline Halo-γ2 signal compared to control treatment (binned LME: time × treatment interaction: F(1, 992) = 1.1088, p = 0.2926). n ≥ 48 puncta per timepoint (control), 24 puncta (Sema4D). (iii) Sema4D treatment increases mean Halo-γ2 intensity colocalized with GFP-Gephyrin puncta in the bottom quintile of baseline Halo-γ2 intensity compared to control treatment (binned LME: time × treatment interaction: F(1, 992) = 4.2854, *p < 0.05). n ≥ 49 puncta per timepoint (control), 24 puncta (Sema4D). Error bars = SEM. Data are normalized within treatment condition to the mean of the first 3 minutes. (C) Sema4D treatment enhances the negative correlation between baseline (t=0 min.) Halo-γ2 fluorescence colocalized with GFP-Gephyrin puncta and the change in colocalized Halo-γ2 fluorescence over time. Control: slope CI = (−2.731×10^−5^, -1.203×10^−5^); Sema4D: slope CI = (−6.294×10^−5^, -3.327×10^-5^). n = 276 puncta (control), 138 puncta (Sema4D). Error bars = 95% CI of linear regression fit. (D) Sema4D treatment decreases the mean nearest neighbor distance of GFP-Gephyrin puncta to Halo-γ2 puncta (for puncta with initial nearest neighbor distance ≤ 2 µm) (binned LME: time × treatment interaction : F(1, 2456) = 50.247, ***p < 0.001). n ≥ 131 puncta per timepoint (control), 54 puncta (Sema4D). Error bars = SEM.

Because individual gephyrin scaffolds vary widely in their GABA_A_R content, we next asked whether the Sema4D-dependent increase in colocalized Halo-γ2 fluorescence was driven preferentially by recruitment to scaffolds with initially low receptor levels. We analyzed the mean colocalized Halo-γ2 fluorescence in the subset of GFP-Gephyrin puncta in the top quintile of baseline Halo-γ2 fluorescence vs. GFP-Gephyrin puncta in the bottom quintile of baseline Halo-γ2 fluorescence. Sema4D treatment significantly increased colocalized Halo-γ2 fluorescence at gephyrin scaffolds in the lowest quintile of baseline receptor expression (Fig. 7Bii) while having no effect on gephyrin scaffolds in the top quintile (Fig. 7Biii), suggesting that Sema4D-dependent GABA_A_R recruitment was primarily directed toward postsynaptic specializations that lacked receptors. In neurons treated with Fc control protein there was a slight negative correlation between baseline Halo-γ2 fluorescence colocalized with GFP-Gephyrin puncta and the change in Halo-γ2 fluorescence over time (Fig. 7C). Sema4D treatment enhanced this negative correlation, again suggesting that Sema4D preferentially promotes recruitment of Halo-γ2 receptors to postsynaptic sites with low baseline GABA_A_R density. Additionally, Sema4D treatment reduced the mean nearest neighbor distance between GFP-Gephyrin and Halo-γ2 puncta relative to control (Fig. 7D). Together these results suggest that Sema4D increases Halo-γ2 localization at gephyrin-labeled scaffolds primarily by redistributing existing receptors to scaffolds lacking GABA_A_Rs rather than by altering overall Halo-γ2 expression levels or receptor density at existing postsynaptic sites.

### Sema4D treatment increases recruitment of GABA_A_Rγ2 to gephyrin scaffolds without driving new colocalization events between GABA_A_Rγ2 and GFP-Gephyrin puncta

While most Halo-γ2 puncta were localized to postsynaptic specializations marked by GFP-Gephyrin, a subset of these puncta appeared to move independently of gephyrin. Pre-clustered GABA_A_R assemblies that are not associated with gephyrin have been described previously (Danglot et al., 2003; Jacob et al., 2005; Christie et al., 2006), but their role in synapse formation is unclear. One possibility is that these clusters may act as pre-assembled packets of receptors that can be rapidly recruited to synapses when required. We hypothesized that Sema4D treatment would increase colocalization events between pre-clustered GABA_A_R puncta and gephyrin puncta. Similar to our previous analysis of GAD65 and gephyrin colocalization (Fig. 5), we identified GFP-Gephyrin puncta that were not initially colocalized with a Halo-γ2 puncta but became colocalized during the imaging session and remained stably colocalized for at least 10 minutes. We found that in contrast to GAD65 and gephyrin, in which the majority of new colocalization events involved emergence of a new gephyrin puncta, most new colocalization events between GFP-Gephyrin and Halo-γ2 puncta (∼80% of events) were between pairs of pre-existing protein puncta (Fig. 8A). We also found that a substantial proportion of GFP-Gephyrin puncta (20-25%) colocalized with a previously independent Halo-γ2 puncta (Fig. 8Bi). Surprisingly, however, new colocalization events occurred at a similar frequency in both control and Sema4D-treated neurons (Fig. 8Bii,iii), suggesting that Sema4D does not drive localization of independent GABA_A_R clusters to postsynaptic scaffolds. Thus, the occurrence of new colocalization events between GFP-Gephyrin and Halo-γ2 puncta fails to explain the Sema4D-dependent increase in total Halo-γ2 recruitment.

**Figure 8.**
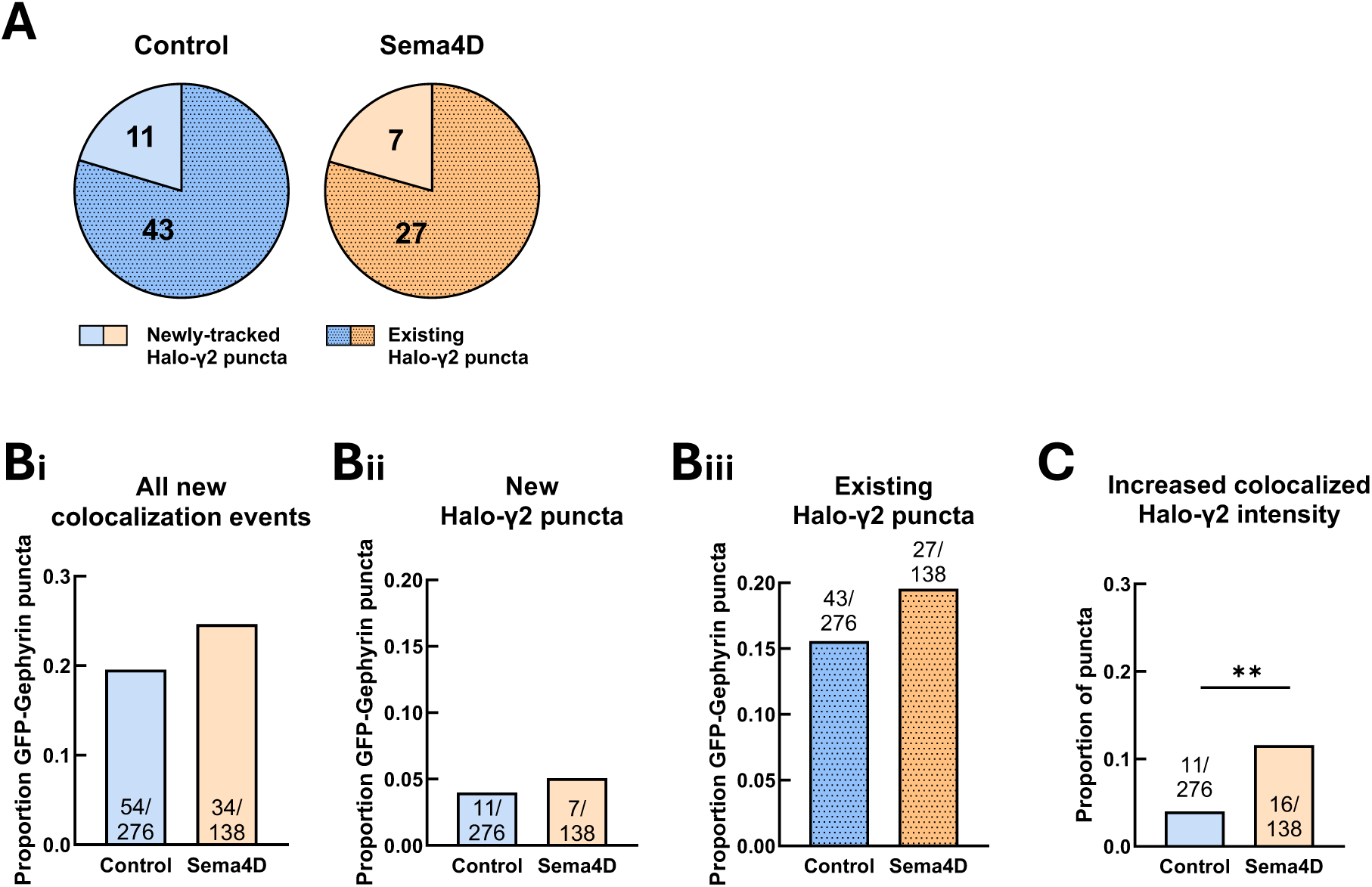
Sema4D treatment increases recruitment of GABA_A_Rγ2 to gephyrin scaffolds without driving new colocalization events between GABA_A_Rγ2 and GFP-Gephyrin puncta. (A) Number of new colocalization events between GFP-Gephyrin and existing Halo-γ2 puncta vs. newly-tracked Halo-γ2 puncta. (B) (i) Overall proportion of GFP-Gephyrin puncta with a new colocalization event. There was no effect of Sema4D treatment on the proportion of GFP-Gephyrin puncta with any type of new colocalization event (p = 0.2525). (ii) There was no effect of Sema4D treatment on the proportion of GFP-Gephyrin puncta that colocalized with a newly-tracked Halo-γ2 puncta (p = 0.6155, Fisher’s exact test). (iii) There was no effect of Sema4D treatment on the proportion of GFP-Gephyrin puncta that colocalized with an existing Halo-γ2 puncta (p = 0.3314). (C) Proportion of GFP-Gephyrin puncta with ≥ 1.5-fold increase in colocalized Halo-γ2 fluorescence. Sema4D treatment significantly increased the proportion of GFP-Gephyrin puncta with increased Halo-γ2 fluorescence (**p < 0.01, Fisher’s exact test).

Similar to our previous analysis of GAD65-GFP, we used principal component analysis (PCA) to determine which GFP-Gephyrin particle tracking parameters most strongly predict new colocalization events with Halo-γ2 puncta. As before, PC1 and PC2 most strongly weighted parameters related to GFP-Gephyrin mobility and proximity to Halo-γ2 puncta, respectively (Table S3). GFP-Gephyrin puncta that colocalized with Halo-γ2 clusters showed a moderate tendency towards decreased mobility along PC1 and decreased nearest neighbor distance to Halo-γ2 puncta along PC2 (Fig. S9A,D). While mean velocity of GFP-Gephyrin puncta that colocalized with Halo-γ2 puncta did not differ from that of other GFP-Gephyrin puncta (Fig. S9B,E), GFP-Gephyrin puncta that colocalized with Halo-γ2 puncta were significantly closer to Halo-γ2 puncta on average (Fig. S9F), suggesting they are located in receptor-rich regions of the dendrite.

### Recruitment of GABA_A_Rs that were not previously associated with GFP-Gephyrin puncta underlies Sema4D-dependent postsynaptic maturation

We identified a third population of GFP-Gephyrin puncta in which colocalized Halo-γ2 fluorescence increased at least 1.5-fold during the imaging session regardless of whether a new colocalization event occurred. We found that, compared to control, Sema4D significantly increased the fraction of GFP-Gephyrin puncta that underwent a >1.5-fold increase in colocalized Halo-γ2 fluorescence from about 4% to 11.5% (Fig. 8C). Next, we examined mobility characteristics for the population of GFP-Gephyrin that underwent a >1.5-fold increase in colocalized Halo-γ2 fluorescence using PCA. Unlike the previous GFP-Gephyrin population that underwent new colocalization events (Fig. S9A-F), these GFP-Gephyrin puncta showed no clear clustering along either PC1 or PC2 (Fig. S9G) and they did not differ from other GFP-Gephyrin puncta or between treatment conditions in their mean velocity (Fig. S9H) or proximity to Halo-γ2 puncta (Fig. S9I). Thus, gephyrin puncta that recruit GABA_A_Rγ2 in response to Sema4D treatment are not necessarily localized to GABA_A_R-rich regions where many receptor clusters are already present. Taken together these results suggest two main routes to receptor localization at postsynaptic scaffolds: 1) colocalization between existing gephyrin scaffolds and pre-formed GABA_A_R clusters in receptor-rich regions, and 2) recruitment of receptors that were not previously clustered to sites where gephyrin is present, with only the latter being regulated by Sema4D treatment.

### New colocalization events between GFP-Gephyrin and Halo-GABA_A_Rγ2 may be established by clusters of either protein

The canonical view posits that relatively stable gephyrin scaffolds recruit mobile GABA_A_Rs to postsynaptic specializations, leading to receptor immobilization and synapse maturation. Colocalization events between gephyrin and pre-clustered GABA_A_R puncta were frequent (Fig. 8B), and while Sema4D treatment did not specifically drive these events, we reasoned that our live imaging approach could nonetheless give us unique insight into this process. We analyzed the paths followed by GFP-Gephyrin and Halo-γ2 puncta before and after new colocalization events, beginning by tracking the total displacement distance of GFP-Gephyrin and Halo-γ2 clusters prior to colocalizing (Fig. 9A, E). In Fc control-treated neurons we observed that, surprisingly, GFP-Gephyrin and Halo-γ2 puncta moved similar distances from their original location prior to colocalization, with few puncta moving more than 2 µm prior to colocalization in all conditions (Fig. 9B, F). These data suggest that either gephyrin or GABA_A_R cluster mobility can promote a new colocalization event. We observed that Sema4D treatment led to a marginally significant increase in the mean distance traveled by GFP-Gephyrin puncta prior to colocalizing (p = 0.0545) (Fig. 9B) and a marginally significant decrease in the average distance of GFP-Gephyrin to colocalization sites in the 10-minute window prior to colocalization (p = 0.0553) (Fig. 9C). This suggests that Sema4D treatment increases mobility of this subset of GFP-Gephyrin puncta earlier than 10 minutes before colocalization with pre-clustered Halo-γ2 puncta. We also observed that GFP-Gephyrin was on average farther from its origin location after colocalization compared to before colocalization in control neurons, but this effect was absent in Sema4D-treated cultures (Fig. 9D) suggesting, in agreement with our analysis of GAD65-GFP and Halo-Gephyrin (Fig. 6), that Sema4D signaling does not direct mobile gephyrin puncta to specific pre-determined locations on the dendrite.

**Figure 9.**
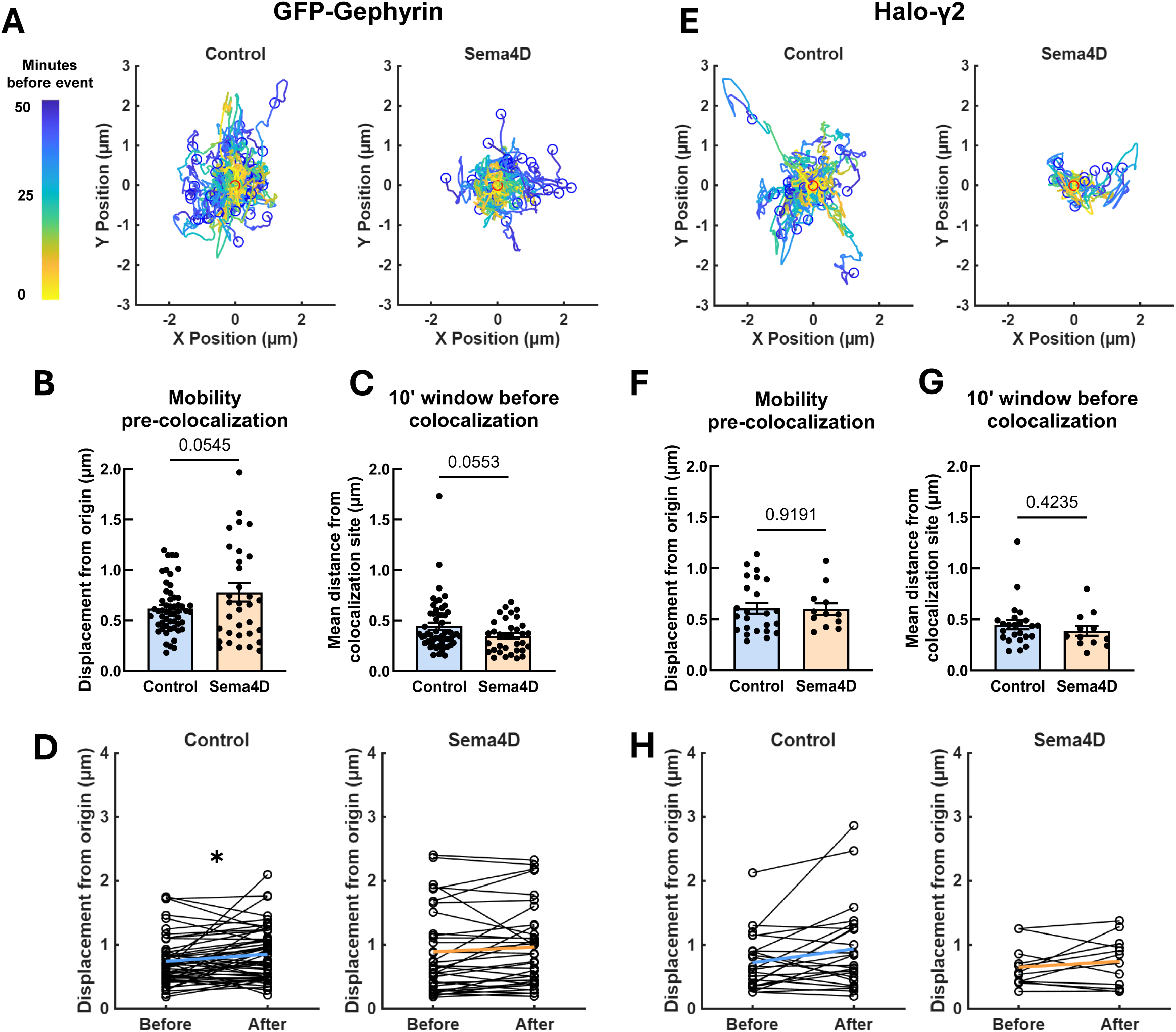
Both GFP-Gephyrin puncta and pre-clustered Halo-γ2 puncta are mobile prior to colocalizing. (A) Tracks followed by GFP-Gephyrin puncta prior to new colocalization events with Halo-γ2 puncta in control (left) or Sema4D (right) treated cultures. Blue dots represent relative start locations; red dots (0,0) represent normalized location of new colocalization event. (B) Displacement distance of GFP-Gephyrin puncta undergoing a new colocalization event prior to colocalizing. Comparison shows displacement from origin of GFP-Gephyrin puncta that colocalized with a Halo-γ2 puncta in Sema4D-treated cultures compared to control (p = 0.0545, Mann-Whitney U-test). (C) Mean distance of GFP-Gephyrin puncta from new colocalization sites in the 10-minute window preceding new colocalization events. Comparison shows mean distance of GFP-Gephyrin puncta undergoing colocalization events with Halo-γ2 puncta in Sema4D-treated cultures compared to control (p = 0.0553). n = 54 events (control), 34 events (Sema4D). Error bars represent SEM. (D) Change in mean displacement from origin of individual GFP-Gephyrin puncta in the 10 min. window before vs. the 10 min. window after colocalization. GFP-Gephyrin puncta are significantly further from origin after colocalization events in control neurons (*p < 0.05, Wilcoxon matched-pairs signed rank test) but not in Sema4D- treated cultures (p = 0.1126). n = 54 events (control), 34 events (Sema4D). (E) Tracks followed by Halo-γ2 puncta prior to new colocalization events in control (left) or Sema4D (right) treated cultures. Blue dots represent relative start locations; red dots (0,0) represent normalized location of new colocalization event. (F) Displacement distance of Halo-γ2 puncta undergoing a new colocalization event with GFP-Gephyrin prior to colocalization. Sema4D treatment does not affect displacement of Halo-γ2 puncta undergoing colocalization events with GFP-Gephyrin puncta (p = 0.9191, Mann-Whitney U-test). (G) Sema4D treatment does not affect mean distance of Halo-γ2 puncta from colocalization sites during the 10-minute window preceding new colocalization events compared to control (p = 0.4235). n = 24 events (control), 12 events (Sema4D). Error bars represent SEM. (H) Change in mean displacement from origin of individual Halo-γ2 puncta in the 10 min. window before vs. the 10 min. window after colocalization. Halo-γ2 puncta are not significantly further from origin after colocalization events in either control (p = 0.0738, Wilcoxon matched-pairs signed rank test) or Sema4D-treated cultures (p = 0.4238). n = 24 events (control), 12 events (Sema4D).

We performed similar analyses on Halo-γ2 mobility before and after colocalization with GFP-Gephyrin and found no difference in any of these parameters between control and Sema4D-treated cultures (Fig. 9F-H). Taken together these data suggest that, contrary to the canonical model, both gephyrin and GABA_A_R protein clusters are mobile and are equally capable of initiating a colocalization event. These new colocalization events appear to occur stochastically and do not require Halo-γ2 puncta to be recruited to pre-established sites. Overall, these experiments suggest a model in which mobile preformed clusters of gephyrin and GABA_A_Rs may encounter each other to form new postsynaptic specializations. While Sema4D modulates gephyrin mobility prior to colocalization, the overall rate of colocalization events is not significantly enhanced; rather, enhanced recruitment or capture of individual GABA_A_Rs by relatively immature postsynaptic scaffolds is the primary mechanism driving Sema4D-mediated receptor recruitment and synapse maturation.

## Discussion

Compared to excitatory synapse formation, inhibitory synapse formation has historically been more challenging to study due to the lack of an extensive postsynaptic density for biochemical purification and the relatively low abundance of inhibitory synapses. The spatiotemporal dynamics of inhibitory synapse assembly are poorly understood, particularly at acute timescales, as the vast majority of assays employed to study the function of synaptogenic molecules have assessed synapse formation by performing a manipulation (e.g. gene knockout) and assaying the presence or absence of synapses by microscopy or electrophysiology. These retrospective approaches likely obscure significant nuances in the spatiotemporal dynamics of synaptic protein cluster formation, stabilization at sites of colocalization, and maturation. Thus there remains a significant gap in understanding the process by which molecular signals and cellular processes transform nascent contacts into mature inhibitory synapses.

Over the past decade we defined a novel role for class 4 Semaphorins and Plexin-B receptors in regulating inhibitory synapse formation in rodent hippocampus. Specifically, our lab demonstrated that the soluble, extracellular domain of Sema4D induces inhibitory synapse formation on a rapid time scale (i.e. minutes) while having no effect on excitatory synapse formation. Among the handful of identified transsynaptic regulators of GABAergic synapse assembly, which include the Neuregulin-ErbB4, Neuroligin-Neurexin, Slitrk-PTPδ, FGF, and Dystroglycan families (Levinson et al., 2005; Krivosheya et al., 2008; Takahashi et al., 2012; Yim et al., 2013; Dabrowski et al., 2015; Trotter et al., 2023), only Sema4D has these unique properties. This rapid, selective, and inducible effect enables precise dissection of synaptogenic mechanisms that are otherwise difficult to study due to the asynchronous and developmentally protracted nature of synaptogenesis.

In this study we leveraged the ability of Sema4D to induce GABAergic synapse formation on the scale of minutes coupled with two-channel live imaging to characterize the dynamics of synaptic proteins during Sema4D-mediated synapse assembly. Our data support a model in which Sema4D treatment increases mobility of presynaptic GAD65-containing protein clusters, allowing GAD65 puncta to explore a wider radius to establish sites of putative new synapses. In contrast, postsynaptic gephyrin-containing protein clusters are mobilized only once they are in the immediate vicinity of a presynaptic bouton where GAD65 is present. Our findings agree with converging evidence from multiple groups that the presynaptic compartment primarily initiates GABAergic synapse formation during early development (Wierenga et al., 2008; Dobie and Craig, 2011; Kuriu et al., 2012). These coordinated dynamics suggest Sema4D primarily drives GABAergic synapse formation by increasing the likelihood of stable colocalization of nearby synaptic protein clusters that are poised to be recruited to existing contacts in the presence of a synaptogenic signal. Only a specific subset of protein clusters that are present near existing contacts appears to be available to form new synapses, and synaptogenic signaling pathways act on these protein clusters to rapidly assemble synapses at acute timescales.

The canonical model of inhibitory postsynaptic assembly which emerged from early single-particle tracking and FRAP experiments posits that gephyrin scaffolds are stable structures which capture laterally diffusing GABA_A_Rs upon synaptic entry (see e.g. Jacob, Bogdanov et al. 2005). Subsequent biochemical work supported this anchoring model, showing that gephyrin and collybistin directly bind GABA_A_Rs to stabilize receptor localization (Tretter et al., 2008; Hines et al., 2018; Lorenz-Guertin and Jacob, 2018). Our live imaging data support this general framework but reveal a more dynamic view of postsynaptic assembly. We found that Sema4D promotes recruitment of GABA_A_Rs to less mature gephyrin scaffolds with an abundance of available binding sites rather than enhancing receptor clustering at sites where many receptors are already present. Notably, Sema4D did not alter the frequency of new colocalization events between pre-assembled gephyrin and GABA_A_Rγ2 puncta (Fig. 8), indicating that the effect of Sema4D is primarily to stabilize or enhance accumulation of individual receptors at postsynaptic sites lacking receptors.

Interestingly, we observed that both gephyrin and GABA_A_Rγ2 could establish new sites of colocalization by moving toward putative new postsynaptic sites, and that this process plays an important role in postsynaptic assembly during development. Recent work suggests that gephyrin turnover and assembly is dynamically regulated by multiple binding (GlyR, GABARAP) and phosphorylation sites (via Erk1/2, GSK-3, Cdk5) (Petrini and Barberis, 2014; Choii and Ko, 2015; Chapdelaine et al., 2021), and that gephyrin assembles into filaments that phase separate to allow for flexible rearrangement of postsynaptic structures rather than forming a rigid lattice as previously believed (Macha et al., 2025). Thus, gephyrin mobility appears to be more dynamic than originally appreciated. These observations challenge the traditional view that receptor clustering is strictly secondary to scaffold formation and raise the possibility that, in some contexts, GABA_A_R clustering may precede or even initiate recruitment of mobile gephyrin assemblies to developing inhibitory synapses.

The molecular mechanisms linking Sema4D signaling to local recruitment of synaptic proteins remain largely unclear. Plexin-B1, the high-affinity ligand of Sema4D, is expressed both pre- and postsynaptically and is required in both compartments for proper recruitment of synaptic components (McDermott et al., 2018; Adel et al., 2024). Whether Sema4D/Plexin-B1 signaling acts simultaneously on both sides of the synapse, or whether signaling from the pre- or postsynaptic compartment acts on the other compartment indirectly, is unknown. Our findings in this study suggest that Sema4D-dependent changes to presynaptic mobility precede localization of gephyrin puncta to new contact sites. This implies that Sema4D likely exerts its presynaptic effects via direct binding to Plexin-B1 expressed in the presynaptic interneuron, while postsynaptic effects are mediated by proximity to an eligible presynaptic bouton. Interestingly, prior work from our lab indicated that knockdown of Plexin-B1 in the postsynaptic neuron prevents Sema4D-dependent synapse formation (McDermott et al., 2018); thus, Plexin-B1 expression in the postsynaptic cell is still required. Sema4D-dependent mobilization of gephyrin puncta once in proximity to a GAD65-positive bouton could presumably be mediated by paracrine interactions between Sema4D and Plexin-B1 or by Plexin-B1 signaling in conjunction with other signaling pathways via coreceptors (Giordano et al., 2002; Swiercz et al., 2004).

Downstream signaling through the intracellular C-terminal GAP and RhoGEF domains of Plexin-B1 mediates cytoskeletal remodeling (Ito et al., 2006; Tran et al., 2007; Vodrazka et al., 2009). One possibility is that Plexin-B1 directly promotes presynaptic protein mobility by regulating microtubule tracks, molecular motors, or force-generating cytoskeletal reorganization (e.g. actin branching or polymerization). A second possibility is that local cytoskeletal disruption or disassembly effectively “releases the brake” on presynaptic bouton mobility, allowing mobile boutons to sample a larger dendritic area, and stabilization subsequently occurs through contact with a postsynaptic specialization, preventing elimination. The latter hypothesis is supported by a study which demonstrated that local application of Sema4D to single boutons induced stabilization which could be chemically mimicked by destabilizing actin filaments (via latrunculin B treatment) or by inhibiting the RhoA/ROCK pathway (Frias et al., 2019). Although further work is required to distinguish between these possibilities, the relatively slow velocities and confined movement radii of protein clusters involved in new synapse formation point to modulation of local actin networks as the main structural change that promotes synapse formation downstream of Plexin-B1 signaling. Together, our data support a model in which Sema4D/Plexin-B1 signaling facilitates presynaptic protein mobility by removing actin-dependent constraints on pre-assembled presynaptic protein clusters and increasing the probability of transient contacts with nearby postsynaptic specializations which are then stabilized through reciprocal adhesion.

Overall, the findings from this study show that Sema4D signaling coordinates dynamic yet locally constrained changes in both pre- and postsynaptic compartments to assemble functional inhibitory synapses on rapid timescales. This capacity to precisely regulate inhibitory synapse formation has important implications for inhibitory circuit organization in the developing and mature brain: inhibitory synapses regulate the timing and synchrony of network activity, and disruptions to genes involved in inhibitory synapse assembly are implicated in various neurodevelopmental and seizure disorders (Shimojima et al., 2011; Sun et al., 2011; Lionel et al., 2013). The ability of Sema4D to coordinate pre- and postsynaptic protein mobility to rapidly assemble new synapses highlights a potential mechanism for fast, stable circuit remodeling and presents an intriguing yet largely unexplored therapeutic angle for disorders of excitatory-inhibitory balance such as epilepsy (Acker et al., 2018; Adel et al., 2023). More broadly, this model provides a general framework for how cellular signaling pathways may tune inhibitory connectivity and circuit balance on behaviorally and clinically relevant timescales.

## Acknowledgements

We would like to thank Dr. Avital Rodal and Dr. Andrew Stone for critical reading of this manuscript, Dr. Adam Puche and Chloe Jenkins at the University of Maryland Medical School for providing GAD65-GFP mice, the Brandeis Light Microscopy Core Facility and Foster Animal Facility for resource access and technical support, and all Paradis Lab members, in particular Susannah Adel, Rabia Anjum, Sarah McCallister, Yi Zhang, and Roshni Ray for critical discussions and experimental design/analysis feedback. This work was supported by NIH grant R01NS065856 (S.P.), a CURE Epilepsy Catalyst Award no. 998742 (S.P.), and NIH fellowship award F31NS134188 (Z.P.).

## Methods

### Ethics statement

All animal procedures were approved by the Brandeis University Institutional Animal Care and Usage Committee, and all experiments were performed in accordance with relevant guidelines and regulations.

### Animals

Male GAD65-GFP transgenic mice (López-Bendito et al., 2004) were obtained courtesy of Dr. Adam Puche (University of Maryland School of Medicine) and maintained in our animal facility with ad libitum access to food and water on a 12 hour day/night cycle. Heterozygous GAD65-GFP males were crossed to female B6CBAF1/J mice (Jackson Laboratories, #100011) to produce litters in which approximately half of pups express one copy of the GAD65-GFP transgene. As this line expresses bright visible green fluorescence throughout the brain and spinal cord, GAD65-GFP pups were identified using a handheld 488 nm laser and an orange emission filter. For rat hippocampal cultures, pregnant female Sprague-Dawley rats were obtained from Charles River Labs when litters were approximately E14 and kept in our facility until dissection.

### Primary mouse hippocampal cultures

For GAD65-GFP and Halo-Gephyrin live imaging experiments, primary rat astrocytes were plated onto 35 mm Petri dishes with glass-bottom 14 mm microwells (MatTek #P35G-1.5-14-C) that had been coated overnight at 37°C with poly-d-lysine (20 μg/ml) and laminin (3.4 μg/ml). Before plating glia, coverslips were washed three times with sterile Ultrapure water and once with DMEM (Gibco #10313039). Glia were plated in DMEM with FBS (GenClone # 25-550) and grown in a 37°C incubator with 5% CO_2_ until confluent. When glia formed a confluent feeder layer, AraC (Sigma #C1768) was added at a final concentration of 5 µM in the dish to prevent further division. At P0–1, GAD65-GFP mouse pups were identified as described above and rapidly sacrificed by decapitation. Hippocampi were harvested from GAD65-GFP pups of both sexes, dissociated with papain (20 units/mL) for 8 minutes, and gently resuspended in Neurobasal medium (Gibco #21103049) with B27 supplement (Gibco #17504044) (NB/B27) before plating atop glia within microwells at a density of 180k cells/well. After 4–24 hrs. when neurons were fully adhered the plating media was replaced by 1.5 mL NB/B27 with 5 µM AraC; the culture media was not changed thereafter.

For GFP-Gephyrin and Halo-GABA_A_Rγ2 overexpression live imaging experiments, astrocytes were grown on 14 mm glass-bottom microwell, 35 mm Petri dishes as described above. Pregnant female Sprague-Dawley rats were sacrificed at embryonic day 18 (E18) by CO_2_ asphyxiation, pups were rapidly removed and decapitated, and heads were kept in ice cold dissociation media prior to dissection. Hippocampi were then dissected from pups of both sexes, dissociated, and resuspended in NB/B27 before plating atop astrocytes at a density of 120k cells/well. After 4–24 hrs. the plating media was replaced by 1.5 mL NB/B27 and was not changed thereafter, except during transfection (see below).

### Infection/transfection and HaloTag labeling

For GAD65-GFP and Halo-Gephyrin live imaging experiments, neurons were infected on DIV2 or DIV3 with AAV9.hSyn-HaloTag-Gephyrin virus (custom-produced by Duke Viral Vector Core) at a final concentration of 1 × 10^9^ GC/mL (∼0.83 × 10^3^ GC/neuron) in the dish. Culture media was not changed after addition of the virus. For labeling of HaloTag-expressing neurons, Janelia Fluor 646 HaloTag Ligand (Promega #GA1110) was suspended in DMSO according to manufacturer recommendations to create a 200 µM stock solution which was aliquoted and stored at -20°C for up to 1 year. To prepare the labeling solution this stock was diluted 1:200 in NB/B27 to create a 1 µM 5x working stock. At DIV10–11 cultures were live labeled by aspirating all but 80 µL of growth media from each dish and adding 20 µL of 5x dye solution for a final concentration of 200 nM dye in the well. Final dilution was such that DMSO comprised no more than 0.1% of the total media volume during labeling. Cultures were incubated with ligand solution at 37°C for 15 minutes. After labeling, cells were briefly washed once with standard NB/B27 media which was then replaced with phenol red-free NB/B27 media prior to imaging. (Note: although excess unbound JF646 HaloTag ligand is reported to be minimally fluorescent, we observed greatly improved signal-to-noise after a brief wash).

For GFP-Gephyrin and Halo-GABA_A_Rγ2 live imaging experiments, DIV4 cultures were transfected with Lipofectamine 2000 using a protocol adapted from (Marwick and Hardingham, 2017). Lipofectamine 2000 reagent (3 µL/ng DNA) was diluted to a final volume of 33 µL per well using NB/B27; plasmid DNA (650 ng total split evenly between the two constructs) was separately diluted to a final volume of 33 µL per well with NB/B27. The L2000 solution was added to the DNA solution, pipetted 5–6x, and left to incubate at RT for 20 minutes. After incubation, the transfection mix was diluted to final volume of 125 µL per well using NB/B27. Working one dish at a time, growth media was aspirated from each dish and quickly replaced with 125 µL diluted transfection mix, and cells were incubated with transfection mix for 2–3 hrs. at 37°C. A recovery media was then made by mixing 80% saved growth media with 20% fresh NB/B27, and transfection media was fully aspirated and replaced with 1.5 mL recovery media. The media was not changed again prior to labeling. At DIV10–11 Janelia Fluor labeling was performed as described above.

### Live imaging

Live images were obtained using an inverted Nikon AX-R Resonance Scanning Confocal with Ti2 body, Nikon Perfect Focus, a piezo Z controller, and a MRD71670 Plan Apochromat Lambda D 1.42 NA 60X Oil objective. Culture dishes were placed into a humidified environmental enclosure maintained at 37°C with 5% CO_2_ at a constant flow rate of 0.2 L/minute. Cells were allowed to habituate for at least 15 minutes prior to imaging. Single GAD65-GFP positive cells were identified and a field of view where distal axons were clearly visible was chosen. Cultures were treated with either with 2 nM human IgG1-Fc control (R&D Systems # 110-HG) or 2 nM recombinant Sema4D-Fc chimera (R&D Systems #7470-S4) by pipetting directly into the dish. Image acquisition was started immediately after adding the protein and setting the focal plane, typically within 1-2 minutes of adding each treatment. For experiments with GAD65-GFP and Halo-Gephyrin, we used 488 and 640 nm laser lines from a LUA-S4 laser unit at 2.5% and 3.0% power respectively, and for experiments with GFP-Gephyrin and Halo-γ2 we used 488 and 640 nm lines with 5.2% and 4.0% power respectively. 12-bit images were acquired at a 2048 × 2048 resolution with a pixel density of 126.6 nm/px using the resonant scanner and 8x line averaging. A Z-stack of 5–7 Nyquist-sampled planes encompassing a total range of 1.5-2.1 µm was acquired at 15 second intervals for 1 hour with optical focusing correction (Nikon Perfect Focus) to minimize drift in the Z direction.

### Image unwarping and registration

To eliminate the possibility that changes to protein cluster mobility were due to cell motility, changes to dendrite/axon morphology, or image drift in the XY plane, images were registered and unwarped prior to particle tracking analysis. Time lapse images were first corrected for any stage XY drift during acquisition using the Linear Stack Alignment with SIFT plugin in ImageJ with the following parameters: initial Gaussian blur 1 px, steps per octave 8, image size 100–250 px, feature descriptor size 4, feature descriptor orientation bins 8, closest/next closest ratio 0.96, maximum alignment error 5 px, inlier ratio 0.05, and rigid transform.

Following linear alignment, images were next unwarped using the BigWarp plugin in ImageJ (Bogovic et al., 2016) to correct for non-translational drift (movement of axons/dendrites, etc.). Briefly, time lapses were flattened into maximum intensity projections and then converted to a virtual Z-stack. The first frame of each image was duplicated repeatedly (once per timepoint) and converted to a virtual Z-stack to form a reference stack that was identical in size to the moving image stack. The moving image stack and reference stack were then imported into BigWarp viewer, and pairs of landmarks were manually chosen in the last plane of the moving image (corresponding to the last timepoint of the time lapse) and the reference stack. To ensure proper linear interpolation of the transformation grid across time, the moving image was split into 30-frame intervals, and landmarks in the moving image were linearly interpolated at the corresponding 30-frame intervals. The 30-frame sub-videos were then individually aligned to the interpolated landmarks using a thin-plate splines transform, which uses a deformable grid to perform exact matching between the moving image and reference image landmarks. Transformations were applied using BigWarp Apply with the following parameters: thin plate spline transformation, bounding type FACES, samples = 5, and linear interpolation. For each 30-frame sub-video, the last frame of the unwarped output served as the reference stack to which the next sub-video was aligned; this prevents compounding error over time. Unwarped 30-frame sub-videos were finally stitched together and converted back to a time series for particle tracking analysis.

### Qualitative scoring of GAD65-GFP protein cluster behavior

Primary hippocampal cultures were generated from P0 GAD65-GFP mice as described above. DIV2 cultures were infected with a virus expressing HaloTag-Gephyrin under the synapsin promoter. Cultures were treated with 2 nM Sema4D-Fc or Fc control protein at DIV11 as previously described and imaged at 10s intervals for 1 hour. To qualitatively assess cellular processes that may be relevant to synapse formation, the following categories of behaviors were chosen based on empirical observation: trafficking, nascent branching, local cluster mobility, splitting, merging, complex split/merge events, active growth cones, stable branch formation, and stable branch removal. Complete descriptions of these behaviors are found in Table S1. An experimenter blinded to condition manually traced 10 axons (including branches) per cell using the freehand line tool in ImageJ. Axons were selected only if the majority of the process remained in focus across all time points. Behaviors were manually counted by an experimenter blinded to condition and the frequency of each behavior was normalized to the total length of each axon.

### Particle tracking for live images

For all analyses of protein cluster mobility and real-time colocalization analysis we utilized the surface tracking feature in Imaris for Neuroscientists v10.2.0 (Oxford Instruments). To facilitate accurate tracking of low-intensity protein clusters, live images were denoised using onboard Nikon Denoise.ai in Nikon Elements prior to generating max intensity projections. Max intensity projections were then imported into Imaris for tracking with both the denoised and raw fluorescence channels. Surfaces were created in the GAD65-GFP, Halo-Gephyrin, GFP-Gephyrin, and/or Halo-γ2 channels using the machine learning segmentation feature in Imaris with the denoised channel; we determined empirically that this led to more consistent identification of protein clusters than standard background subtraction and thresholding, particularly for bright GAD65-GFP clusters along axons. We next tracked surfaces across frames using the following parameters: autoregressive motion tracking, max frame-to-frame distance 2 µm, and max frame gap 8. Tracks were automatically filtered out if their duration was less than 60 seconds and were manually removed if they showed spurious linkages between adjacent neurites or if they were located on a neurite that did not appear to be an axon (for GAD65-GFP) or a dendrite (for gephyrin or Halo-γ2). For most analyses, only protein clusters that could be tracked for the duration of the live imaging session were considered for further analysis.

### Analysis of protein cluster mobility

#### Surface and track features

All analyses were performed in MATLAB R2025a (MathWorks) and visualized in Graphpad Prism 10.5.0. Surface and track statistics for each channel were exported from Imaris and fed into a custom MATLAB analysis pipeline for data workup. Surface features that were exported for analysis included surface area, acceleration, displacement delta length, intensity in each channel, position, overlapped area ratio, speed, and nearest neighbor distance to other surfaces. Track features that were exported for analysis included track duration, straightness, AR(1) mean, length, and number of surfaces. Other track features were calculated within our analysis pipeline, such as raw change in fluorescence (F_end_ - F_0_), fold change fluorescence (F_end_ / F_0_), peak fold change fluorescence (F_max_ / F_0_), and Euclidean distance (net displacement between selected timepoints).

#### Fluorescence intensity analysis

All analysis of fluorescence intensity was performed using the raw fluorescence (i.e., non-denoised) channel. For analysis of protein cluster fluorescence intensity over time, a photobleach correction was applied by fitting a one-step exponential decay model to the average particle intensity in the control condition using 𝐹(𝑡) = 𝐴 ⋅ 𝑒^−𝑘𝑡^ + 𝐶, where A is the amplitude, k is the decay rate constant, t is time, and C is the baseline offset. To correct photobleaching the raw fluorescence was transformed using 𝐹_𝑐𝑜𝑟𝑟𝑒𝑐𝑡𝑒𝑑_(𝑡_𝑛_) = (𝐹_𝑚𝑒𝑎𝑠𝑢𝑟𝑒𝑑_(𝑡_𝑛_) ⋅ 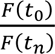). We then measured the average and total fluorescence intensity within the outline of each individual puncta for each timepoint. For analysis of active GAD65-GFP regions (Fig. S4), we quantified the total combined fluorescence intensity within GAD65-GFP puncta within a band drawn with a 1 µm radius around the GAD65-GFP puncta. For figures measuring the change in normalized fluorescence intensity over time, fluorescence intensity was normalized to the baseline mean of the first three minutes (12 frames) for all puncta within each treatment condition.

#### Principal component analysis

Dimensionality reduction was performed using principal component analysis (PCA) via MATLAB’s default pca function. Track statistics for key parameters of puncta size, intensity, mobility, and proximity were calculated and tabulated as shown in Table S2–3. All track parameters were z-score normalized before PCA. All tracks were then plotted along PC1 and PC2, and function outputs for component weights and percent track variance explained were saved. Track parameters were considered significant contributors to a PC if the absolute value of their weighting for a PC exceeded 0.4.

#### New colocalization event analysis

New colocalization events were defined as instances in which a GAD65-GFP puncta that was not colocalized with a gephyrin puncta (or vice versa) in the previous frame became colocalized, which we determined by comparing the nearest neighbor distance to a Halo-Gephyrin puncta between frames. To avoid counting transient crossings, both puncta were required to remain colocalized in at least 95% of frames over the next 10 minutes following the initial colocalization event (colocalization events happening in the final 10 minutes of the imaging session were not analyzed). To determine whether the puncta in the opposite channel was newly-tracked (new puncta) or previously present (existing puncta), we identified puncta in the opposite channel for which the center of mass was located within 1 µm of the colocalization site at the colocalization timepoint, and used this criterion to determine the unique track ID of the opposite-channel puncta. If the opposite-channel puncta was present at least 10 minutes before the colocalization event, it was considered an existing puncta; otherwise it was classified as a new puncta. After classifying new colocalization events, we then quantified average protein cluster velocity, fluorescence intensity, displacement from origin, and distance from the colocalization site in 1) the entire window before the new colocalization event and 2) the 10-minute periods before and after the new colocalization event.

### Experimental design and statistical analysis

Statistical analyses for all figure panels except time series data were performed using Graphpad Prism 10.5.0. For each experiment, specific statistical tests are described in the figure legends. Data were tested for normality before appropriate statistical tests were applied. Where indicated, outliers were identified by the ROUT method with Q = 1% and removed from the data set.

For time series data, to assess whether Sema4D treatment interacts with GAD65-GFP puncta features to regulate gephyrin accumulation in GAD65-GFP puncta, we fit binned time series data to linear mixed-effects models (LME) using MATLAB (fitlme), which was chosen to account for the effects of cell-to-cell variability on track measurements. Due to the large number of timepoints sampled in each image, timepoints were binned to avoid artificially inflating statistical power, with bin size determined by calculating autocorrelation at a range of bin sizes for the mean of the control condition across timepoints using MATLAB autocorr function. Bin size was fixed as the smallest round-number bin size at which the autocorrelation function dropped below 0.3, thus ensuring semi-independence of timepoints for the purposes of LME model fitting. Models tested the effects of treatment condition, time, GAD65-GFP puncta displacement length, area, and/or intensity, and all 2-way and 3-way interactions, on either: the nearest distance between GAD65 and gephyrin puncta, or mean fluorescence intensity of gephyrin within GAD65-positive regions. Random intercepts were used to account for variation between images (within-experiment variability) and between individual tracks within images (within-puncta variability over time). Model fit was assessed by inspecting residuals, random effects distributions, and summary fit indices (AIC, BIC, LogLikelihood). Effect estimates are reported as changes in the dependent variable per unit change in the predictor. Time-dependent effects are interpreted as rate changes per minute. For main effects and interactions with p < 0.05, follow-up analysis was conducted using subsets of puncta according to the variable of interest (e.g., a significant interaction between time, treatment, and puncta size was further analyzed by grouping puncta by quintiles according to size and plotting mean gephyrin fluorescence over time in each subset.) For experiments involving GFP-Gephyrin and Halo-γ2, statistical analysis of live imaging parameters was performed identically to the above.

## Code availability

All ImageJ and Python scripts used for landmark interpolation, unwarping, and stitching are publicly accessible via GitHub at zpranske/Bigwarp_Analysis. All MATLAB scripts and functions used for statistical analysis of particle tracking data are publicly accessible via GitHub at zpranske/LiveImaging_Analysis_Imaris.

