## Supplementary Information for "Coordinated pre- and postsynaptic protein dynamics underlie rapid Sema4D-mediated inhibitory synapse assembly"

**Fig. S1**

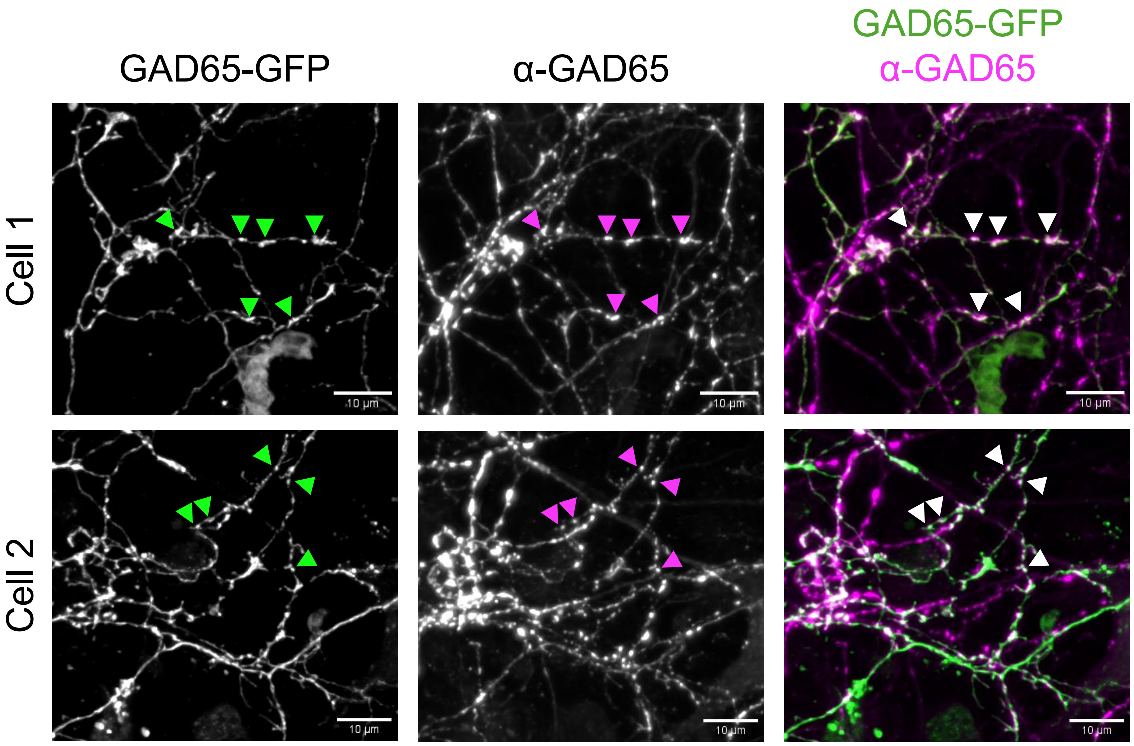

**Figure S1. Representative images of DIV11 mouse hippocampal neurons expressing GAD65-GFP.**

GAD65-GFP labeled puncta reliably colocalize with GAD65 antibody staining in distal axons of primary cultured DIV11 hippocampal neurons from GAD65-GFP mice. Green arrows = GAD65-GFP; magenta arrows = anti-GAD65. The majority of GAD65-GFP puncta in GFP-positive cells are marked by anti-GAD65 antibody (white arrows = colocalized). Scale bars = 10 µm.

Fig. S2

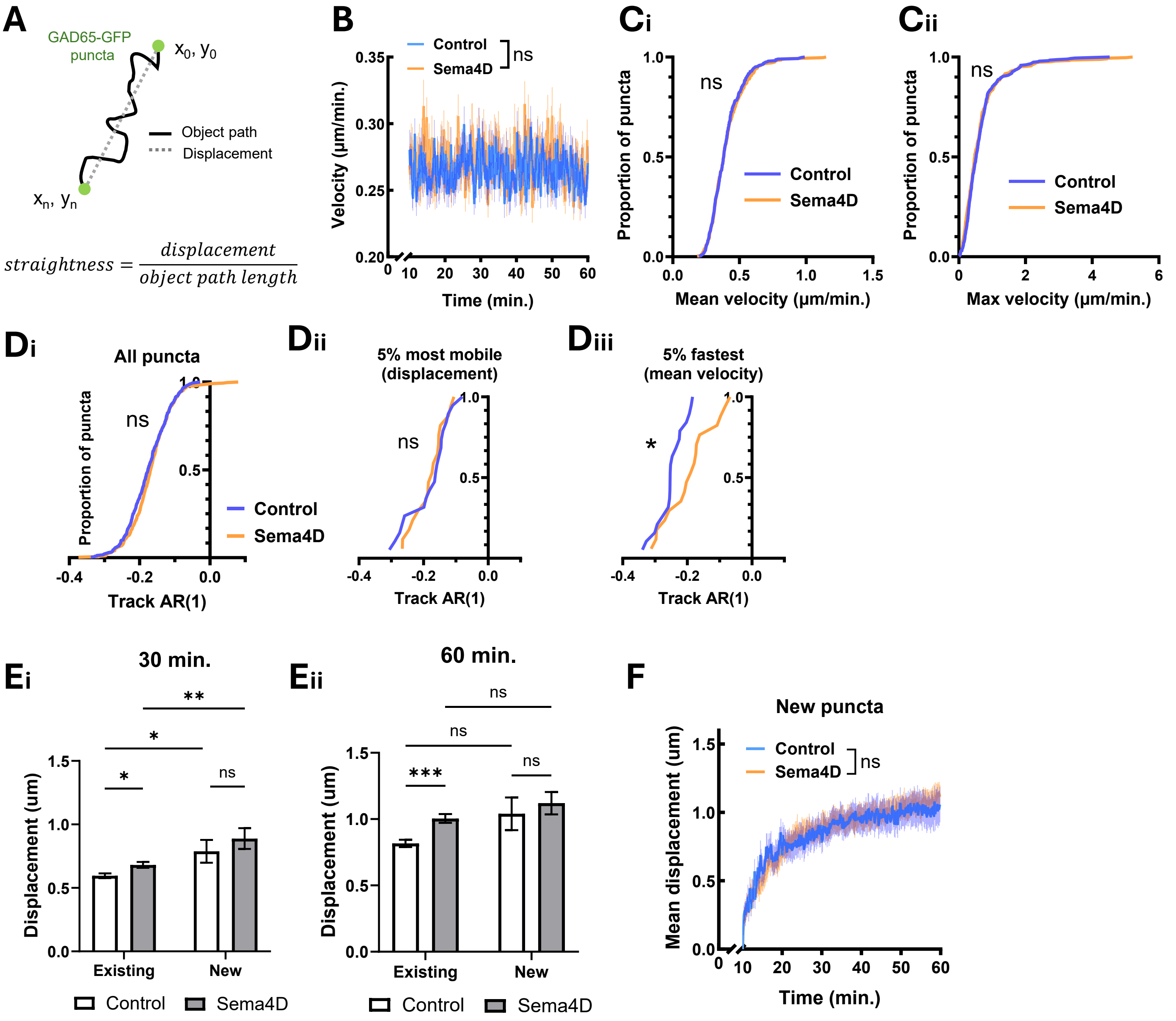
A
GAD65-GFP puncta
$x_0, y_0$
$x_n, y_n$
Object path
Displacement
$straightness = \frac{displacement}{object\ path\ length}$
B
Control
Sema4D
ns
Velocity (μm/min.)
Time (min.)
Ci
ns
Control
Sema4D
Proportion of puncta
Mean velocity (μm/min.)
Cii
ns
Control
Sema4D
Proportion of puncta
Max velocity (μm/min.)
Di
All puncta
ns
Control
Sema4D
Proportion of puncta
Track AR(1)
Dii
5% most mobile (displacement)
ns
Track AR(1)
Diii
5% fastest (mean velocity)
*
Track AR(1)
Ei
30 min.
**
*
*
ns
Displacement (um)
Existing
New
Control
Sema4D
Eii
60 min.
ns
ns
***
ns
Displacement (um)
Existing
New
Control
Sema4D
F
New puncta
Control
Sema4D
ns
Mean displacement (um)
Time (min.)

**Figure S2. Additional data related to GAD65-GFP mobility.**

(A) Graphic indicating definitions of puncta displacement and track straightness.

(B) Mean velocity of GAD65-GFP puncta over time. Sema4D treatment does not significantly alter mean puncta velocity (binned LME: time × treatment interaction: $F(1, 7306) = 0.0268$ , $p = 0.870$).

(C) (i) 1h Sema4D treatment did not affect the distribution of GAD65-GFP puncta mean velocity (Kolmogorov-Smirnov test, $p = 0.9542$) or (ii) GAD65-GFP puncta maximum velocity ($p = 0.4656$).

(D) Autoregressive parameter (AR(1)) for GAD65-GFP puncta tracks in each condition. (i) There was no significant difference between Sema4D and control for the entire population of GAD65-GFP puncta (Kolmogorov-Smirnov test, $p = 0.1416$) or for (ii) the top 5% most mobile puncta by displacement ($p = 0.3561$). (iii) The top 5% fastest puncta by mean velocity had significantly higher AR(1) parameters in Sema4D-treated cultures compared to control (*$p < 0.05$), indicating that these puncta followed more directionally consistent paths.

(E) Displacement of existing and newly-tracked (appearing between 3-20') GAD65-GFP puncta at 30' and 60'. At 30' newly-tracked puncta are more mobile than existing puncta (Two-way ANOVA: $p < 0.0001$). Post hoc test showed this effect was present in both control (*$p < 0.05$, Holm-Šídák multiple comparisons test) and Sema4D-treated cultures (**$p < 0.01$). Although at 60' there was a main effect of new vs. existing puncta on mobility (Two-way ANOVA: $p < 0.01$), this effect was not significant in post hoc tests for control ($p = 0.0807$) or Sema4D (0.5088).

(F) Displacement of newly-tracked GAD65-GFP puncta in control vs. Sema4D-treated cultures. Sema4D did not affect mobility of newly-tracked puncta (binned LME: time × treatment interaction: $F(1, 1631) = 0.5091$ , $p = 0.4756$). n = 68 puncta (Fc), 102 puncta (Sema4D). Note: analysis begins at t=10 min.

Fig. S3

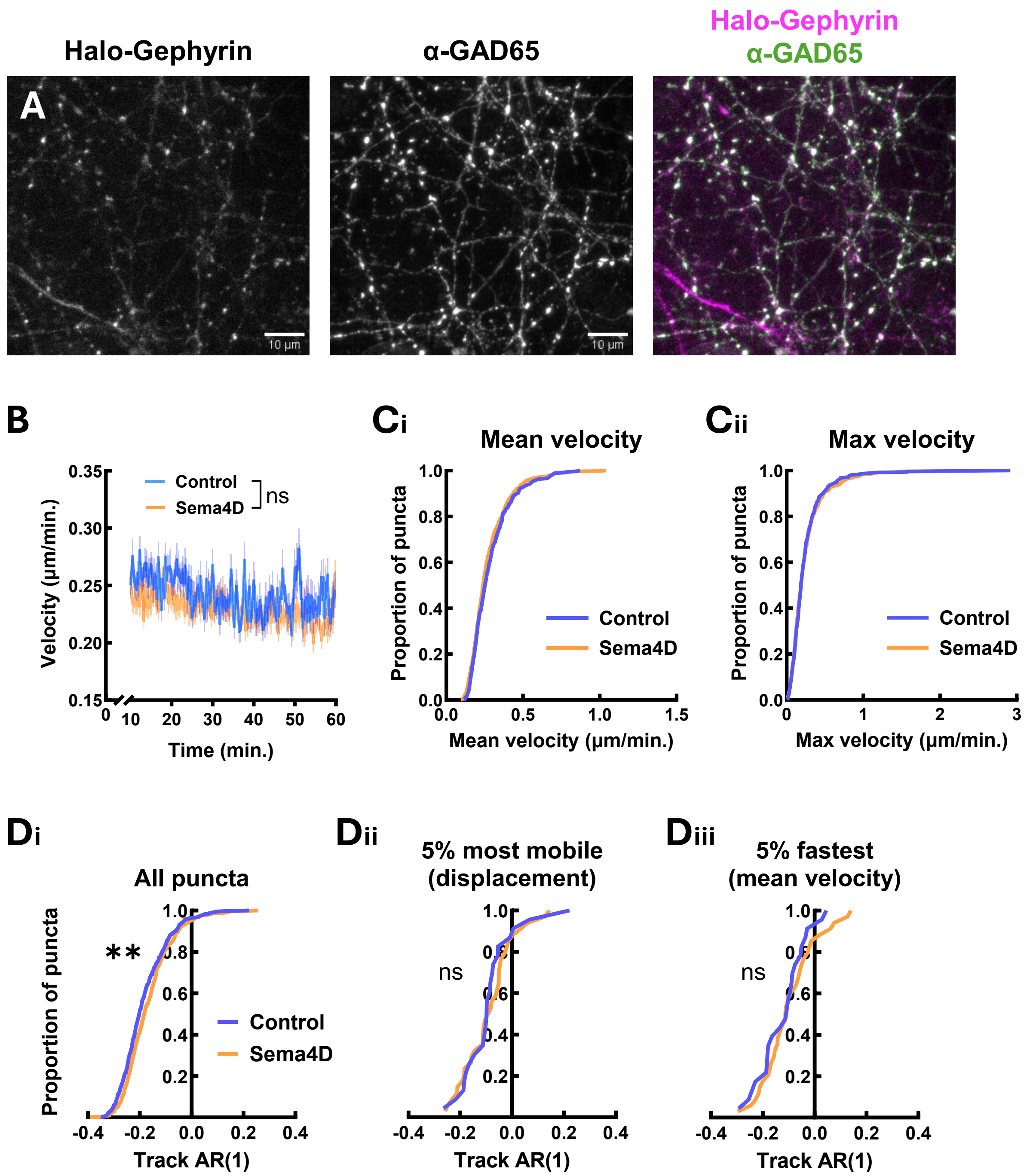

Halo-Gephyrin
α-GAD65
Halo-Gephyrin
α-GAD65
A
10 µm
10 µm
B
Control
Sema4D
ns
Velocity (µm/min.)
Time (min.)
Ci
Mean velocity
Proportion of puncta
Control
Sema4D
Mean velocity (µm/min.)
Cii
Max velocity
Proportion of puncta
Control
Sema4D
Max velocity (µm/min.)
Di
All puncta
Proportion of puncta
**
Control
Sema4D
Track AR(1)
Dii
5% most mobile
(displacement)
ns
Track AR(1)
Diii
5% fastest
(mean velocity)
ns
Track AR(1)

**Fig. S3. Additional data related to Halo-Gephyrin mobility.**

(A) Virally-expressed Halo-Gephyrin in cultured E18 rat neurons colocalizes with anti-GAD65 antibody. Magenta = Halo-Gephyrin, green = anti-GAD65 antibody, white = colocalized. Scale bar = 10 μm.

(B) Mean velocity of Halo-Gephyrin puncta over time. Sema4D treatment does not significantly alter mean puncta velocity (binned LME: time × treatment interaction: $F(1, 9042) = 0.3542$ , $p = 0. 5518$).

(C) (i) 1h Sema4D treatment did not affect the distribution of Halo-Gephyrin puncta mean velocity (Kolmogorov-Smirnov test, $p = 0.1919$) or (ii) Halo-Gephyrin maximum velocity ($p = 0.8541$).

(D) Autoregressive parameter (AR(1)) for Halo-Gephyrin puncta tracks in each condition. (i) Sema4D significantly shifted the distribution of Halo-Gephyrin track AR(1) parameters for the entire population of Halo-Gephyrin puncta (Kolmogorov-Smirnov test, $p < 0.01$). However, there was no difference for (ii) the top 5% most mobile puncta by displacement (center; $p = 0.5902$) or (iii) the top 5% fastest puncta by mean velocity (right; $p = 0.9234$).

**Fig. S4**

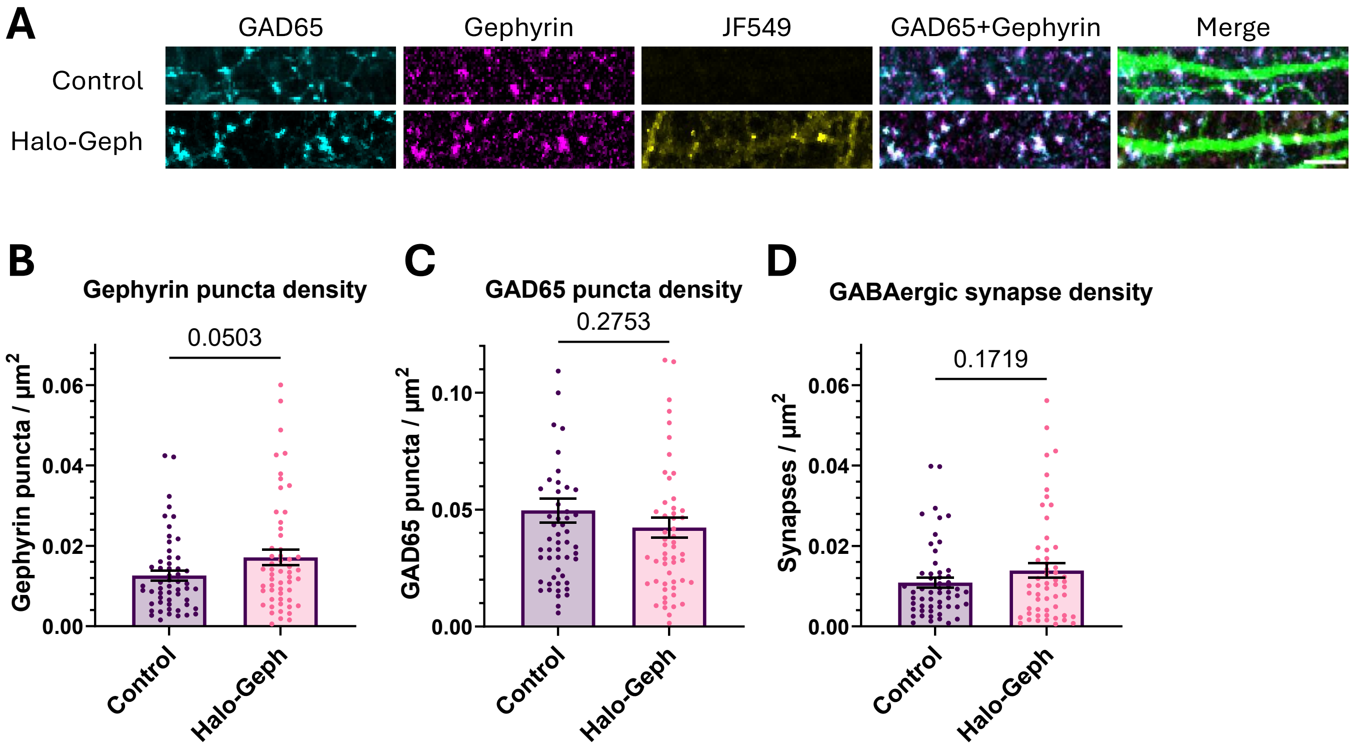
A
GAD65
Gephyrin
JF549
GAD65+Gephyrin
Merge
Control
Halo-Geph
B
Gephyrin puncta density
0.0503
Gephyrin puncta / μm2
0.06
0.04
0.02
0.00
Control
Halo-Geph
C
GAD65 puncta density
0.2753
GAD65 puncta / μm2
0.10
0.05
0.00
Control
Halo-Geph
D
GABAergic synapse density
0.1719
Synapses / μm2
0.06
0.04
0.02
0.00
Control
Halo-Geph

**Figure S4. Viral expression of Halo-Gephyrin does not increase GABAergic synapse density.**

(A) Sample stretches of dendrite from DIV11 rat neurons expressing GFP (green) with or without Halo-Gephyrin. Scale bar = 5 µm.

(B) Viral Halo-Gephyrin expression marginally increases gephyrin puncta density compared to no-virus control neurons (unpaired t-test, $p = 0.0503$). $n = 55$ neurons from 2 replicates per condition.

(C) Viral Halo-Gephyrin expression does not affect GAD65 puncta density compared to no-virus control neurons (unpaired t-test, $p = 0.2753$). $n = 55$ neurons from 2 replicates per condition.

(D) Viral Halo-Gephyrin expression does not affect colocalized GAD65/gephyrin synapse density compared to no-virus control neurons (unpaired t-test, $p = 0.1719$). $n = 55$ neurons from 2 replicates per condition.

**Fig. S5**

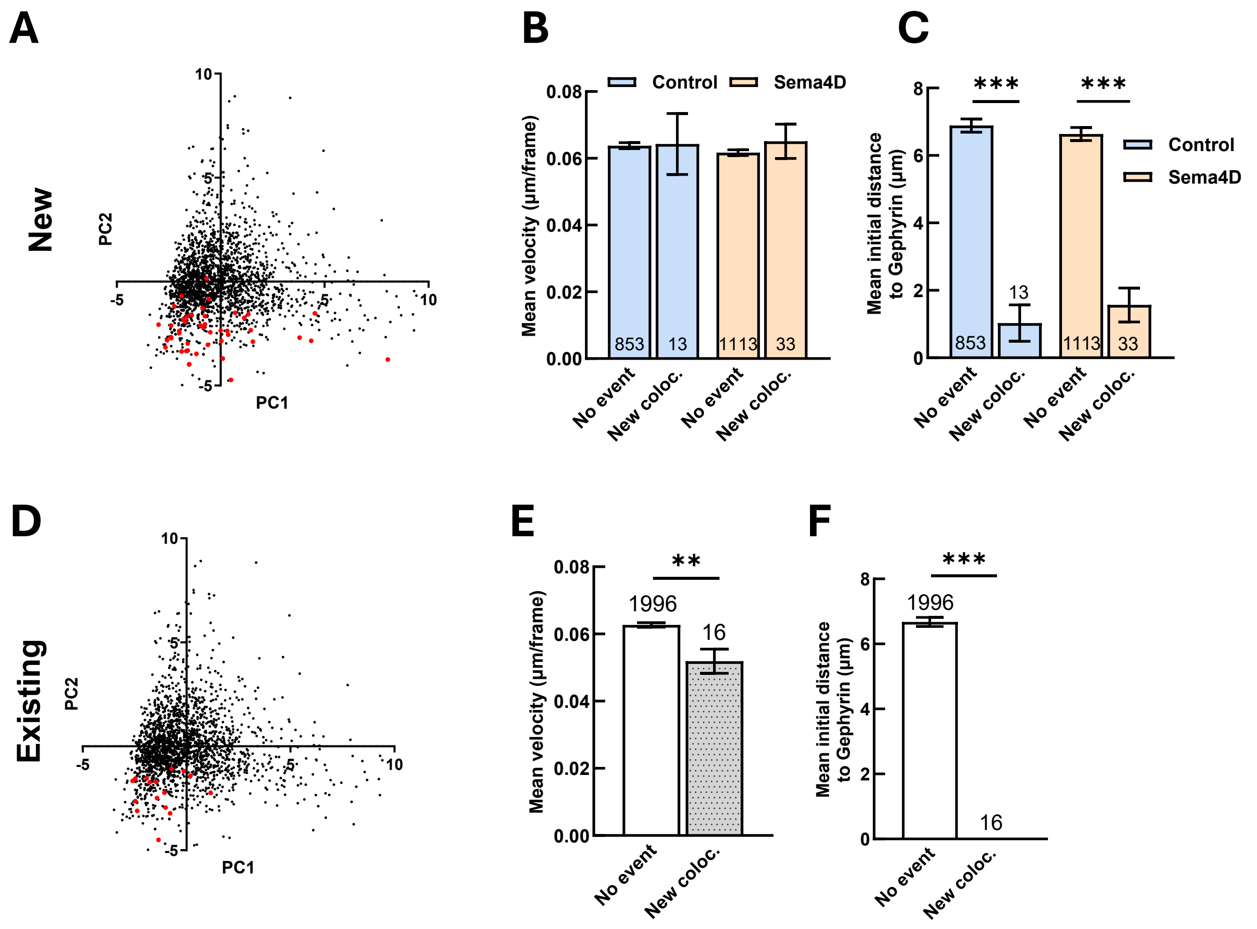

A
B
C
New
PC2
PC1
Control
Sema4D
Mean velocity (µm/frame)
Mean initial distance to Gephyrin (µm)
***
***
853
13
1113
33
No event
New coloc.
D
E
F
Existing
**
1996
16

**Fig. S5. GAD65-GFP puncta that undergo new colocalization events show distinctive profiles of mobility and proximity to gephyrin puncta.**

(A) Scatter plot of GAD65-GFP puncta for principal components (PC) 1 and 2. Red dots = GAD65-GFP puncta that colocalized with a newly-tracked Halo-Gephyrin puncta (n = 46 puncta); black dots = all other GAD65-GFP puncta (n = 1966).

(B) Mean velocity of GAD65-GFP puncta that colocalized with a newly-tracked Halo-Gephyrin puncta does not differ from mean velocity of other GAD65-GFP puncta in control neurons ($p = 0.960$, unpaired heteroscedastic t-test) or in Sema4D-treated neurons ($p = 0.510$). n = number of puncta per condition; error bars = SEM.

(C) Initial distance to nearest gephyrin neighbor is smaller for GAD65-GFP puncta that colocalized with a newly-tracked Halo-Gephyrin puncta compared to other GAD65-GFP puncta in both control neurons (***$p < 0.001$, unpaired heteroscedastic t-test) and in Sema4D-treated neurons (***$p < 0.001$). There was no effect of Sema4D treatment for GAD65-GFP puncta that colocalized with a new gephyrin puncta in control vs. Sema4D treated cultures ($p = 0.469$). n = number of puncta per condition; error bars = SEM.

(D) Scatter plot of GAD65-GFP puncta for principal components (PC) 1 and 2. Red dots = GAD65-GFP puncta that colocalized with an existing Halo-Gephyrin puncta (n = 16 puncta); black dots = all other GAD65-GFP puncta (n = 1996).

(E) Mean GAD65-GFP puncta velocity was significantly decreased for GAD65-GFP puncta that colocalized with an existing Halo-Gephyrin puncta compared to GAD65-GFP puncta without a new colocalization event (**$p < 0.01$). n = number of puncta per condition; error bars = SEM.

(F) Initial distance to nearest gephyrin neighbor is smaller for GAD65-GFP puncta that colocalized with an existing Halo-Gephyrin puncta compared to GAD65-GFP puncta without a new colocalization event (***$p < 0.001$, unpaired heteroscedastic t-test). n = number of puncta per condition; error bars = SEM.

Fig. S6

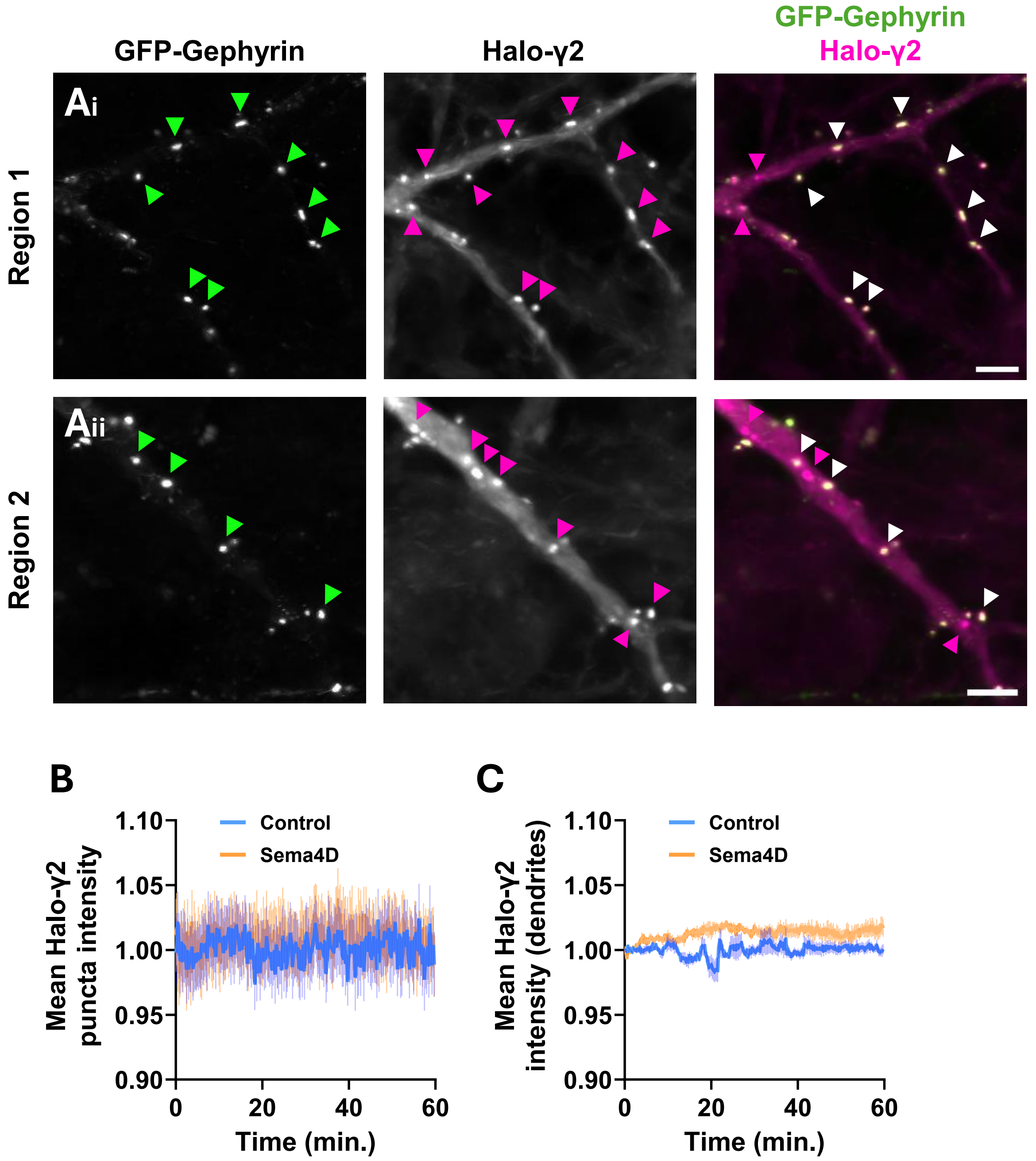
GFP-Gephyrin
Halo-γ2
GFP-Gephyrin
Halo-γ2
Ai
Aii
Region 1
Region 2
B
C
Mean Halo-γ2 puncta intensity
Mean Halo-γ2 intensity (dendrites)
Control
Sema4D
1.10
1.05
1.00
0.95
0.90
0
20
40
60
Time (min.)

**Figure S6. Sema4D does not affect mean Halo-γ2 puncta intensity or dendritic intensity.**

(A) Representative regions showing expression pattern of GFP-Gephyrin (green) and Halo-γ2 (magenta) along dendrites of cultured E18 rat neurons. Most Halo-γ2 clusters are colocalized with GFP-Gephyrin (white arrows), but some independent clusters are observed (magenta arrows). Note diffuse extrasynaptic expression of Halo-γ2 along dendrite. Images represent different regions from the same neuron. Scale bar = 5 μm.

(B) There was no effect of Sema4D treatment on mean intensity of Halo-γ2 puncta (binned LME: time × treatment interaction: $F(1, 1508) = 0.0017$ , $p = 0.9670$). $n \geq 69$ puncta per timepoint (control), 45 puncta (Sema4D). Error bars = SEM. Data are normalized within treatment condition to the mean of the first 3 minutes.

(C) There was no effect of Sema4D treatment on mean total dendritic expression of Halo-γ2 (binned LME: time × treatment interaction: $F(1, 140) = 1.3144$ , $p = 0.2536$). n = 7 cells (control), 5 cells (Sema4D). Error bars = SEM.

**Fig. S7**

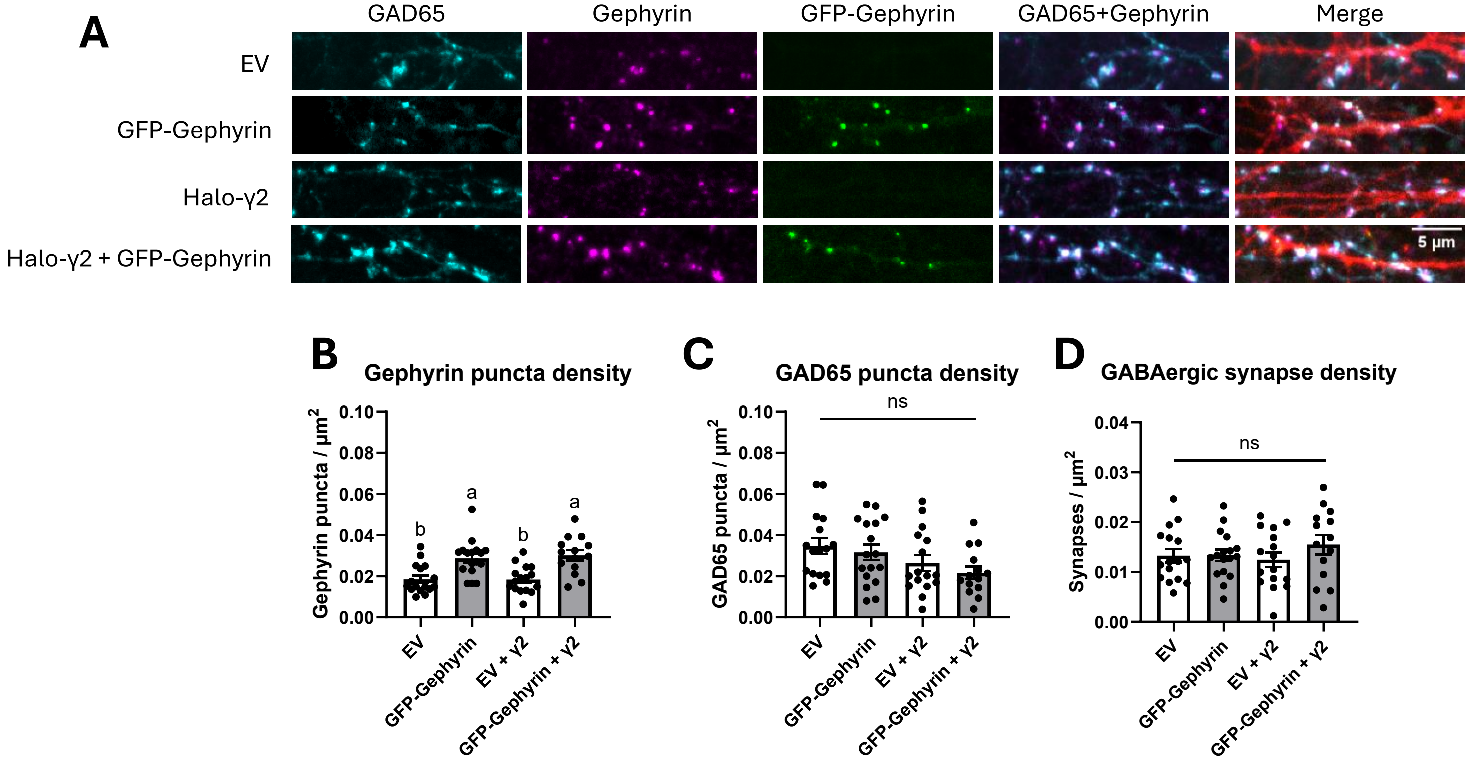
A
GAD65
Gephyrin
GFP-Gephyrin
GAD65+Gephyrin
Merge
EV
GFP-Gephyrin
Halo-γ2
Halo-γ2 + GFP-Gephyrin
5 µm
B
Gephyrin puncta density
Gephyrin puncta / µm2
0.10
0.08
0.06
0.04
0.02
0.00
b
a
b
a
EV
GFP-Gephyrin
EV + γ2
GFP-Gephyrin + γ2
C
GAD65 puncta density
ns
GAD65 puncta / µm2
0.10
0.08
0.06
0.04
0.02
0.00
EV
GFP-Gephyrin
EV + γ2
GFP-Gephyrin + γ2
D
GABAergic synapse density
ns
Synapses / µm2
0.04
0.03
0.02
0.01
0.00
EV
GFP-Gephyrin
EV + γ2
GFP-Gephyrin + γ2

**Figure S7. Overexpression of GFP-Gephyrin increases gephyrin puncta density but does not affect GABAergic synapse density; Halo-γ2 overexpression does not affect gephyrin puncta or synapse density.**

(A) Sample stretches of dendrite from neurons co-expressing TdTomato (red) with indicated constructs. Scale bar = 5 μm.

(B) GFP-Gephyrin overexpression (OE) increases gephyrin puncta density on cultured neurons compared to empty vector (EV) control transfection ($p = 0.0027$, Tukey post-hoc), whereas Halo-γ2 OE does not affect gephyrin puncta density ($p > 0.99$). Co-expression of GFP-Gephyrin and Halo-γ2 does not increase gephyrin puncta density more than GFP-Gephyrin alone ($p = 0.95$).

(C) Overexpression of GFP-Gephyrin alone, Halo-γ2 alone, or combined overexpression of GFP-Gephyrin + Halo-γ2 does not affect density of GAD65 puncta ($F(3, 59) = 2.256$, $p = 0.0912$).

(D) Overexpression of GFP-Gephyrin alone, Halo-γ2 alone, or combined overexpression does not affect GAD65/gephyrin synapse density compared to EV control transfection ($F(3, 59) = 1.676$, $p = 0.536$, ordinary one-way ANOVA).

### Fig. S8

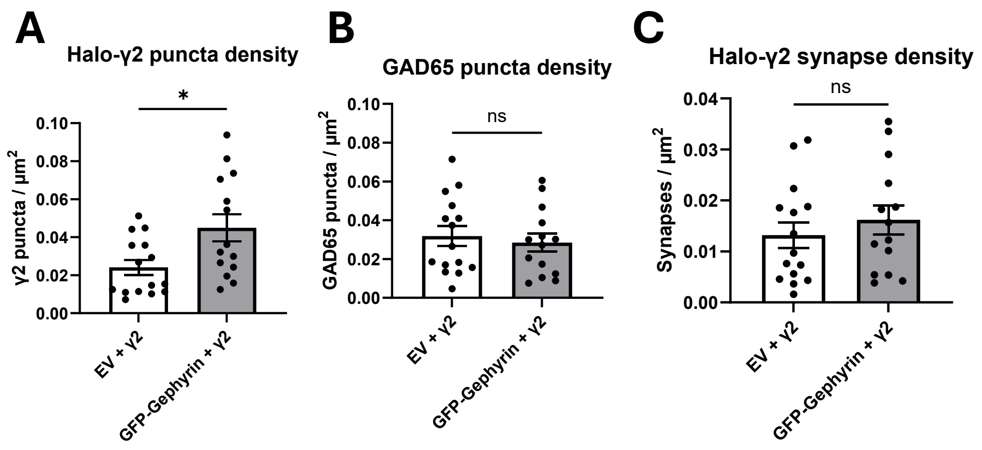
A
Halo-γ2 puncta density
*
γ2 puncta / μm2
0.10
0.08
0.06
0.04
0.02
0.00
EV + γ2
GFP-Gephyrin + γ2
B
GAD65 puncta density
ns
GAD65 puncta / μm2
0.10
0.08
0.06
0.04
0.02
0.00
EV + γ2
GFP-Gephyrin + γ2
C
Halo-γ2 synapse density
ns
Synapses / μm2
0.04
0.03
0.02
0.01
0.00
EV + γ2
GFP-Gephyrin + γ2

**Figure S8. GFP-Gephyrin overexpression does not affect synaptic localization of Halo-γ2.**

(A) Co-expression of GFP-Gephyrin + Halo-γ2 and increases the density of Halo-γ2 compared to Halo-γ2 OE alone ($p = 0.0143$, unpaired t-test).

(B) GFP-Gephyrin OE does not affect density of GAD65+ inputs to transfected cells ($p = 0.6309$).

(C) Co-expression of GFP-Gephyrin and Halo-γ2 does not affect Halo-γ2+ synapse density compared to Halo-γ2 OE alone ($p = 0.436$).

Fig. S9

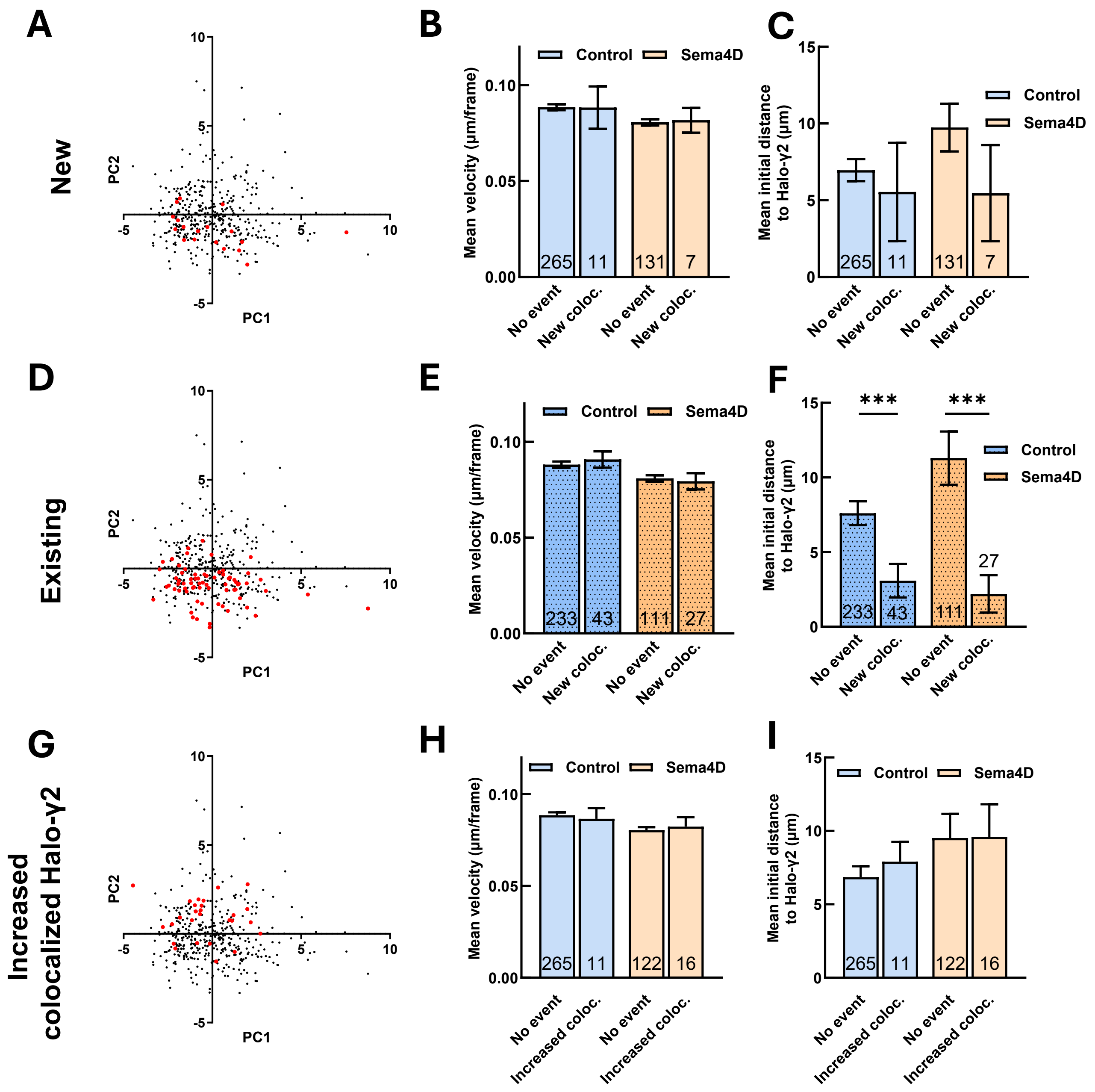

A
New
PC2
PC1
B
Control
Sema4D
Mean velocity (µm/frame)
0.10
0.05
0.00
265
11
131
7
No event
New coloc.
No event
New coloc.
C
Mean initial distance to Halo-γ2 (µm)
15
10
5
0
Control
Sema4D
265
11
131
7
No event
New coloc.
No event
New coloc.
D
Existing
PC2
PC1
E
Control
Sema4D
Mean velocity (µm/frame)
233
43
111
27
No event
New coloc.
No event
New coloc.
F
***
***
Control
Sema4D
Mean initial distance to Halo-γ2 (µm)
233
43
111
27
No event
New coloc.
No event
New coloc.
G
Increased colocalized Halo-γ2
PC2
PC1
H
Control
Sema4D
Mean velocity (µm/frame)
265
11
122
16
No event
Increased coloc.
No event
Increased coloc.
I
Control
Sema4D
Mean initial distance to Halo-γ2 (µm)
265
11
122
16
No event
Increased coloc.
No event
Increased coloc.

**Fig. S9. Principal component analysis of particle tracking parameters of GFP-Gephyrin puncta.**

(A) Scatter plot of GFP-Gephyrin puncta for principal components (PC) 1 and 2. Red dots = puncta with a new colocalization event with a newly-tracked Halo-γ2 puncta (n = 18 puncta); black dots = all other puncta (n = 396).

(B) Mean velocity of GFP-Gephyrin puncta that colocalized with a newly-tracked Halo-γ2 puncta does not differ from mean velocity of GFP-Gephyrin without a colocalization event in control neurons ($p = 0.987$, unpaired heteroscedastic t-test) or in Sema4D-treated neurons ($p = 0.875$). n = number of puncta per condition; error bars = SEM.

(C) Initial nearest neighbor distance to Halo-γ2 puncta for GFP-Gephyrin puncta that colocalized with a newly-tracked Halo-γ2 puncta compared to GFP-Gephyrin puncta without a colocalization event. Puncta that colocalized with a newly-tracked Halo-γ2 puncta did not differ from other GFP-Gephyrin puncta in either control ($p = 0.675$, unpaired heteroscedastic t-test) or Sema4D-treated neurons ($p = 0.251$). n = number of puncta per condition; error bars = SEM.

(D) Scatter plot of GFP-Gephyrin puncta for principal components (PC) 1 and 2. Red dots = puncta that colocalized with an existing Halo-γ2 puncta (n = 70 puncta); black dots = all other puncta (n = 344).

(E) Mean velocity of GFP-Gephyrin puncta that colocalized with an existing Halo-γ2 puncta does not differ from mean velocity of GFP-Gephyrin without a colocalization event in control neurons ($p = 0.553$, unpaired heteroscedastic t-test) or in Sema4D-treated neurons ($p = 0.744$). n = number of puncta per condition; error bars = SEM.

(F) Initial nearest neighbor distance to Halo-γ2 puncta is significantly decreased for GFP-Gephyrin puncta with a new colocalization event with an existing Halo-γ2 puncta compared to other GFP-Gephyrin puncta in both control (***$p < 0.001$, unpaired heteroscedastic t-test) and Sema4D-treated neurons (***$p < 0.001$). However, initial nearest neighbor difference for GFP-Gephyrin puncta with a new colocalization event did not differ between control and Sema4D treatment ($p = 0.599$). n = number of puncta per condition; error bars = SEM.

(G) Scatter plot of GFP-Gephyrin puncta for principal components (PC) 1 and 2. Red dots = puncta with a ≥ 1.5-fold increase in Halo-γ2 fluorescence (n = 27 puncta); black dots = all other puncta (n = 387).

(H) Mean velocity of GFP-Gephyrin puncta with a ≥ 1.5-fold increase in Halo-γ2 fluorescence does not differ from mean velocity of other GFP-Gephyrin puncta in control neurons ($p = 0.760$, unpaired heteroscedastic t-test) or in Sema4D-treated neurons ($p = 0.727$). n = number of puncta per condition; error bars = SEM.

(I) Initial nearest neighbor distance to Halo-γ2 puncta is not significantly different for GFP-Gephyrin puncta with increased Halo-γ2 fluorescence compared to other GFP-Gephyrin puncta in control neurons ($p = 0.503$, unpaired heteroscedastic t-test) or in Sema4D-treated neurons ($p = 0.144$). n = number of puncta per condition; error bars = SEM.

**Table S1. Descriptions of qualitative scoring parameters**

| Category/ behavior | Description |
|---|---|
| Trafficking | Rapid movement of distinct protein clusters outside a radius of ~10-20 µm. Defined as mobility that can only be accomplished by active transport via molecular motors based on speed and long distances covered. This includes only discretely recognizable single clusters. "Flow" events, where the quantity of total fluorescence fluctuates rapidly in a region with apparent directionality, were not counted, although these events were closely associated with trafficking events and are probably due to trafficking of multiple protein clusters that are too numerous/dim/small to be distinguished. Directionality was not considered. |
| Nascent branching | Formation of nascent (non-stable) axonal branches. Each site with branch(es) originating from it was counted as one nascent branch, even if multiple nascent branches were formed and removed over the imaging session. Does not include nascent branching at growth cones, which was categorized separately. |
| Local mobility | Local mobility, defined as movement of a protein cluster within ~5-10 µm of its original location which does not appear to be associated with trafficking. |
| Split | Cluster undergoing a single split event into two or more clusters. Each newly-split cluster is defined as a separate split event. Excludes repeated/complex split/merge events, although a single split and merge was counted. |
| Merge | Merging of two or more clusters into a single cluster. If more than two clusters merge, each merge is counted separately. Excludes repeated/complex split/merge events, although a single split and merge was counted. For practical reasons we required that the merged cluster stop at the merge site for several frames, to ensure that the clusters truly merged (rather than temporarily moving close enough to be indistinguishable) and that a trafficked protein cluster was not just transiently passing. |
| Complex split/merge | Complex repeated split/merge events. Defined as events where two or more nearby protein clusters repeatedly split/merge such that it is difficult to identify individual split/merge events. These usually included 3+ boutons and 2+ split/merge events and were often accompanied by local mobility of one or more of the clusters involved. |
| Growth cone | Presence of a growth cone, characterized by its morphology and distinct movement pattern. Each growth cone was counted even if it later collapsed. |
| Stable branch | Formation of a new stable branch that was at least several microns long and did not appear to be in the process of being removed by the end of the image. Does not include lengthening of existing branches. |
| Branch removed | Removal of an existing axonal branch or collapse of a growth cone. |

### Table S2. GAD65-GFP track principal components and coefficient weights

| | 1 | 2 | 3 | 4 | 5 | 6 | 7 | 8 | 9 | 10 | 11 | 12 | 13 | 14 | 15 |
|---|---|---|---|---|---|---|---|---|---|---|---|---|---|---|---|
| total_distance_traveled | 0.491181 | -0.16826 | -0.14312 | 0.039887 | 0.005215 | -0.04392 | -0.01822 | -0.17292 | 0.059806 | -0.00337 | -0.02888 | -0.10709 | 0.029949 | -0.22257 | 0.780922 |
| mean_net_displacement | 0.241208 | -0.06122 | 0.515127 | 0.258803 | -0.22996 | -0.11529 | -0.10007 | 0.048871 | 0.140749 | -0.00227 | 0.070661 | 0.698157 | 0.009408 | -0.11882 | -0.02684 |
| mean_velocity | 0.498073 | -0.15111 | -0.07683 | 0.061018 | 0.039784 | 0.01438 | 0.054266 | -0.22108 | 0.014951 | -0.00232 | -0.00673 | 0.013929 | -0.07527 | 0.791782 | -0.18111 |
| max_velocity | 0.25184 | -0.06791 | -0.04574 | 0.030857 | 0.001791 | 0.21276 | 0.588094 | 0.731266 | -0.04656 | -0.01331 | 0.020234 | 0.002883 | 0.000909 | -0.01617 | 0.004622 |
| mean_acceleration | 0.480084 | -0.18155 | -0.17594 | 0.028744 | -0.02534 | -0.07436 | -0.05478 | -0.12805 | 0.103343 | -0.00884 | -0.03853 | -0.21041 | 0.031443 | -0.51708 | -0.59525 |
| track_straightness | 0.034295 | -0.00044 | 0.60507 | 0.247219 | -0.2371 | -0.09844 | -0.11996 | 0.154053 | 0.092341 | 0.005309 | -0.07974 | -0.66475 | -0.00955 | 0.098463 | 0.031234 |
| track_ar1 | 0.05214 | 0.118742 | 0.369077 | 0.11308 | 0.220029 | 0.358973 | 0.417313 | -0.4637 | -0.49129 | -0.021 | -0.01146 | -0.0254 | 0.006217 | -0.15551 | -0.0305 |
| mean_area | 0.195416 | -0.07145 | 0.133668 | -0.11544 | 0.489082 | -0.07634 | -0.52367 | 0.346678 | -0.53066 | -0.06061 | -0.00431 | 0.026995 | 0.001919 | 0.002888 | -0.00625 |
| initial_shortest_distance_to_geph | 0.125865 | 0.45993 | -0.11892 | 0.253293 | 0.062013 | 0.008091 | -0.07482 | 0.058525 | -0.01924 | 0.788702 | -0.24105 | 0.02738 | 0.007919 | -0.0046 | -0.00545 |
| min_shortest_distance_to_geph | 0.098718 | 0.494951 | -0.12962 | 0.235606 | 0.011797 | -0.00165 | -0.04776 | 0.040689 | 0.044893 | -0.58285 | -0.56884 | 0.059354 | 0.002115 | 0.00739 | -0.00034 |
| mean_shortest_distance_to_geph | 0.137219 | 0.505695 | -0.11869 | 0.244039 | 0.034369 | -0.03919 | -0.06031 | 0.018419 | 0.01786 | -0.18137 | 0.77645 | -0.09451 | -0.00773 | -0.00495 | 0.001159 |
| gad65_mean_intensity | -0.04524 | -0.04269 | 0.184925 | 0.069167 | 0.70802 | 0.226173 | -0.00023 | -0.00723 | 0.635652 | -0.00214 | 0.005972 | -0.01139 | 0.005205 | -0.02001 | 7.14E-05 |
| gephyrin_mean_intensity | -0.17118 | -0.29323 | -0.20031 | 0.551864 | -0.0102 | 0.205838 | -0.12475 | 0.029461 | -0.09149 | -0.01746 | 0.009021 | 0.004795 | -0.68478 | -0.05722 | 0.012723 |
| gephyrin_f0 | -0.18177 | -0.29547 | -0.18772 | 0.57467 | 0.060975 | -0.00453 | -0.02026 | 0.009318 | -0.11091 | -0.02383 | 0.014863 | -0.00037 | 0.701989 | 0.062522 | -0.01238 |
| gephyrin_df_f | 0.08771 | 0.03137 | -0.02066 | -0.14104 | -0.30125 | 0.833216 | -0.38418 | 0.048162 | 0.067999 | -0.0063 | 0.028854 | 0.00339 | 0.174259 | 0.014773 | -0.00059 |

| pc | % variance explained |
|---|---|
| 1 | 23.1951 |
| 2 | 19.4998 |
| 3 | 13.7128 |
| 4 | 10.8063 |
| 5 | 8.2355 |
| 6 | 6.6855 |
| 7 | 5.587 |
| 8 | 5.2783 |
| 9 | 3.9406 |
| 10 | 1.8892 |
| 11 | 0.493 |
| 12 | 0.4242 |
| 13 | 0.1269 |
| 14 | 0.0865 |
| 15 | 0.0392 |

### Table S3. GFP-Gephyrin track principal components and coefficient weights

| | 1 | 2 | 3 | 4 | 5 | 6 | 7 | 8 | 9 | 10 | 11 | 12 | 13 | 14 | 15 |
|---|---|---|---|---|---|---|---|---|---|---|---|---|---|---|---|
| total_distance_traveled | 0.4973 | -0.0454 | -0.1311 | -0.0458 | -0.1013 | -0.0914 | -0.1233 | -0.1797 | -0.0283 | -0.0106 | 0.0246 | -0.0519 | 0.0235 | -0.1752 | 0.7936 |
| mean_net_displacement | 0.2166 | 0.0108 | 0.5607 | 0.291 | -0.0011 | -0.1945 | -0.0736 | -0.0744 | 0.0061 | -0.0226 | 0.0013 | 0.6925 | -0.1508 | -0.032 | -0.034 |
| mean_velocity | 0.5006 | -0.0577 | -0.0814 | -0.0196 | -0.1374 | 0.0558 | -0.0264 | -0.2205 | -0.0324 | -0.0112 | 0.043 | -0.064 | 0.0192 | 0.7787 | -0.2322 |
| max_velocity | 0.3805 | 0.0332 | -0.049 | 0.1222 | -0.0677 | 0.1507 | 0.1766 | 0.8636 | 0.1278 | 0.0177 | -0.1217 | 0.0162 | -0.0288 | -0.0007 | 0.0045 |
| mean_acceleration | 0.4839 | -0.0344 | -0.1545 | -0.0688 | -0.0794 | -0.1896 | -0.1801 | -0.1316 | -0.0312 | -0.01 | 0.0407 | -0.1365 | 0.0015 | -0.557 | -0.5589 |
| track_straightness | 0.0519 | 0.0336 | 0.6201 | 0.3179 | 0.0458 | -0.1852 | -0.0264 | 0.0196 | 0.0151 | 0.0423 | -0.0118 | -0.6712 | 0.1375 | 0.0213 | 0.0332 |
| track_ar1 | 0.1241 | -0.0558 | 0.1722 | 0.1317 | -0.1775 | 0.7292 | 0.4791 | -0.2954 | 0.0065 | 0.0241 | 0.0436 | -0.0197 | 0.0023 | -0.2254 | -0.028 |
| mean_area | -0.1025 | -0.1507 | -0.2879 | 0.5795 | -0.1929 | -0.0001 | -0.1408 | -0.1235 | -0.0634 | 0.1313 | -0.6696 | -0.0158 | -0.0762 | -0.0013 | -0.0119 |
| initial_shortest_distance_to_y2 | 0.0521 | 0.476 | -0.1422 | 0.1529 | 0.1761 | -0.0396 | 0.0683 | -0.1851 | 0.7881 | -0.1682 | -0.0652 | 0.0009 | 0.0439 | 0.0059 | -0.0055 |
| min_shortest_distance_to_y2 | 0.0613 | 0.4593 | -0.1116 | 0.2253 | 0.2413 | 0.0638 | 0.0489 | 0.0043 | -0.5328 | -0.6099 | -0.0222 | -0.016 | 0.0274 | -0.0021 | 0.0003 |
| mean_shortest_distance_to_y2 | 0.0931 | 0.4494 | -0.1129 | 0.1378 | 0.3356 | 0.0577 | 0.0047 | -0.034 | -0.2291 | 0.749 | 0.1388 | 0.019 | -0.0855 | 0.0086 | 0.0024 |
| geph_mean_intensity | -0.1109 | -0.2075 | -0.2661 | 0.5798 | -0.1297 | -0.018 | -0.1016 | 0.0649 | 0.0692 | -0.0493 | 0.7043 | 0.0086 | 0.0506 | 0.0007 | 0.0066 |
| y2_mean_intensity | 0.0836 | -0.3652 | -0.0866 | 0.0592 | 0.4904 | -0.1753 | 0.3677 | -0.0645 | 0.0407 | -0.0854 | 0.0065 | -0.133 | -0.641 | 0.0142 | 0.0143 |
| y2_f0 | 0.1104 | -0.3683 | -0.0555 | 0.075 | 0.5636 | 0.0139 | 0.1047 | -0.0239 | 0.0092 | 0.0272 | -0.1028 | 0.1484 | 0.6929 | -0.0114 | -0.0201 |
| y2_df_f | -0.0319 | 0.1168 | -0.1081 | 0.0217 | -0.3352 | -0.5346 | 0.7082 | -0.0559 | -0.1121 | 0.0864 | 0.0034 | 0.0561 | 0.2171 | 0.0006 | -0.0025 |

| pc | % variance explained |
|---|---|
| 1 | 24.2555 |
| 2 | 20.0683 |
| 3 | 12.9695 |
| 4 | 12.0055 |
| 5 | 10.6424 |
| 6 | 6.8155 |
| 7 | 6.0191 |
| 8 | 3.3479 |
| 9 | 1.5928 |
| 10 | 1.2286 |
| 11 | 0.5678 |
| 12 | 0.2465 |
| 13 | 0.1652 |
| 14 | 0.062 |
| 15 | 0.0133 |
